# Performance in even a simple perceptual task depends on mouse secondary visual areas

**DOI:** 10.1101/2020.06.25.171884

**Authors:** Hannah C Goldbach, Bradley Akitake, Caitlin E Leedy, Mark H Histed

**Author notes:** For correspondence (M.H.H.). These authors contributed equally to this work (Hannah C Goldbach, Bradley Akitake).

## Abstract

Primary visual cortex (V1) in the mouse projects to numerous brain areas, including several secondary visual areas, frontal cortex, and basal ganglia. While it has been demonstrated that optogenetic silencing of V1 strongly impairs visually-guided behavior, it is not known which downstream areas are required for visual behaviors. Here we trained mice to perform a contrast-increment change detection task, for which substantial stimulus information is present in V1. Optogenetic silencing of visual responses in secondary visual areas revealed that their activity is required for even this simple visual task. *In vivo* electrophysiology showed that, although inhibiting secondary visual areas could produce some feedback effects in V1, the principal effect was profound suppression at the location of the optogenetic light. The results show that pathways through secondary visual areas are necessary for even simple visual behaviors.

## Introduction

Several kinds of neural computations are required to perform sensorimotor tasks. For example, it has been thought that some cerebral cortical neurons represent aspects of the sensory world in neural activity, and other neurons decode that representation into a form useful for motor action. An important step in understanding brain function is determining where representations are present, and where computations like decoding occur. These questions depend on knowing which areas are involved during sensorimotor behaviors. Here, we ask whether secondary visual areas are involved in a simple visual behavior, or whether such behavior can be performed by operating on (“decoding”) sensory representations in V1 or other areas directly, bypassing secondary areas.

The rodent visual system includes several different cortical visual areas, which all respond to visual stimuli, but emphasize different aspects of representation. The borders and locations of mouse visual areas were initially identified through reversals in retinotopy (***Dräger, 1975***; ***Wagor et al., 1980***) and cytoarchitecture (***Caviness, 1975***). Later, injections of anterograde tracers into V1 showed that V1 projects to several secondary visual areas, or higher visual areas, of the mouse brain (***Wang and Burkhalter, 2007***). More recently, the boundaries of cortical visual areas in mice have been examined through mapping retinotopy via intrinsic imaging (***Schuett et al., 2002; Kalatsky and Stryker, 2003; Garrett et al., 2014; Juavinett et al., 2017***) and calcium imaging (***Andermann et al., 2011; Murakami et al., 2017; Zhuang et al., 2017***). Imaging studies have identified as many as sixteen individual, retinotopically mapped areas (***Zhuang et al., 2017***) with differing strengths of connectivity to V1. The most heavily studied secondary visual areas, which receive the largest proportions of projections from V1, are the lateral and medial output pathway areas: LM (lateromedial), AL (anterolateral), RL (retrolateral), and PM (posteriomedial) (***Froudarakis et al., 2019***). While more than half of V1 projections target these four secondary visual areas (***Han et al., 2018***), V1 also projects to a number of other brain areas including frontal cortex and basal ganglia (***Han et al., 2018; Khibnik et al., 2014***), both of which are involved in decision-making and action selection.

We sought to determine whether activity in higher visual areas was essential for a simple visual task. Alternatively, it could be that such activity is not always used, and for some tasks, information flows out of V1 via projections that bypass secondary areas (such as V1’s projections to frontal cortex or basal ganglia). As an example, consider an analogous question in a larger species. In the primate visual object (“where”) pathway, V1 carries representations of small visual edges (***Hubel and Wiesel, 1968***), while areas in inferotemporal (IT) cortex represent complex objects such as faces (***Tsao, 2014***). Identifying the orientation of contours and edges could be carried out by a neural computation that reads out, or decodes, V1 directly, bypassing secondary visual areas such as those in IT cortex. If V1 is read out directly, one might expect that suppression of IT cortical areas, without affecting V1, would impact tasks that require face perception, but leave tasks that involve orientation discrimination unaffected. On the other hand, some data (e.g. on unconscious vs. conscious perception, ***Leopold, 2012***) supports the idea that macaque V1 is simply a conduit for information to secondary visual areas, with no behaviorally relevant direct readout. In the mouse, we are able to investigate which cortical areas are involved in a given behavior using transgenic lines expressing optogenetic proteins, which allow the inactivation of several different cortical visual areas just by moving an optogenetic light spot. To study how secondary visual areas contribute to mouse perceptual behavior, we chose to study a basic task in which V1 contains substantial information. We trained animals to report changes in stimulus contrast.

Neural activity in V1 carries substantial information about visual stimulus contrast (***Albrecht and Hamilton, 1982***; ***Sclar et al., 1990; Vaiceliunaite et al., 2013***). Neurons in mouse V1 are tuned for oriented visual stimuli, with a tuning curve that reaches a peak at the cell’s preferred orientation. The same neurons simultaneously encode contrast: as stimulus contrast is increased, single V1 neurons increase their firing rates monotonically (***Busse et al., 2011; Glickfeld et al., 2013; Khastkhodaei et al., 2016***). As expected from single neurons’ responses, the V1 neural population response varies monotonically with stimulus contrast, and thus the average or summed V1 response does not completely saturate, even up to 100% contrast (***Glickfeld et al., 2013; Histed, 2018***). These features of the population response suggest that mouse V1 activity can be simply (linearly) decoded to allow animals to detect a change in contrast of a visual stimulus. While the structure of population neural variability could in principle make linear decoding diffcult, linear decoders that do not take trial-to-trial variability into account perform well on decoding orientation from mouse V1 (***Berens et al., 2012; Rumyantsev et al., 2020; Stringer et al., 2019; Kafashan et al., 2020***). This suggests that linear decoders would also be able to extract stimulus contrast from the V1 population. Thus, a contrast change detection task does not seem to require any information in more complex representations beyond V1. If the mouse brain had circuitry to decode V1 directly, secondary visual areas might not be involved in the task and inhibiting them would not impact animals’ behavior.

To study whether secondary visual areas are involved in contrast change detection, mice were trained on a contrast-increment detection task in which they report the presence of a flashed Gabor by releasing a lever. This task was previously found to require V1; that is, suppressing V1 optogenetically produced retinotopically-specific impairments in contrast-detection performance (***Glickfeld et al., 2013; Jin and Glickfeld, 2020***). Through optogenetically stimulating all inhibitory neural classes (via the *VGAT-ChR2-EFYP* transgenic mouse line; ***Zhao et al., 2011***) we first confirm this finding: inhibition of V1 negatively affects performance in this task. We then inhibit secondary visual areas, and find that suppression of lateral areas (e.g. LM/AL, and RL) and medial areas (PM) also negatively affect performance. Via electrophysiology, we find that inhibition of secondary visual areas primarily acts to suppress secondary area visual responses, rather than suppressing V1 responses via feedback. Thus, these data show that secondary visual areas are involved in a task that, in principle, could have been performed by the mouse reading out V1 directly. We also find that the effect size caused by secondary area suppression can be large, as suppressing lateral visual areas (LM/AL and RL) produces behavioral effects of similar or even higher magnitude than V1 at the same optogenetic light intensity. In contrast, suppression of PM, the largest medial area, only weakly affected the animals’ performance. Paralleling these behavioral findings, any feedback effects on V1 neural activity were negligible when PM was suppressed, but could be seen in LM at high light intensity, arguing that LM and V1 activity is linked to some extent. Finally, the behavioral effects in all areas are primarily due to sensitivity (*d*′) and not due to changes in false alarm rates. In sum, lateral secondary visual areas (LM/AL and RL) form an interconnected network with V1 that contributes to performance in even simple visually-guided behaviors, where substantial information is available in, and linearly decodable from, V1.

## Results

We studied mice of both sexes expressing ChR2 in all inhibitory neurons (*VGAT-ChR2-EYFP* mouse line, ***Zhao et al., 2011***). We implanted a cranial window and identified visual cortical areas using hemodynamic intrinsic imaging (Methods, ***Figure 1, Figure 1–Figure Supplement 1***). We cemented a fiber-optic cannula over the cranial window to create a light spot targeted to the area of visual cortex we wished to inhibit, and trained these animals on a contrast-increment change detection task (Methods, ***Figure 1A***). Behaving animals were head-fixed in front of an LED monitor and trained to release a lever in response to the onset (increase in contrast, from a neutral gray background) of a Gabor patch for a liquid reward. We varied the contrast change magnitude in order to obtain psychometric curves, which measure the contrast needed for animals to perform the task at a set percentage correct — their perceptual threshold (*e.g*. ***Figure 1C***, open circles; below we discuss effects also on perceptual sensitivity, or *d*′).

**Figure 1.**
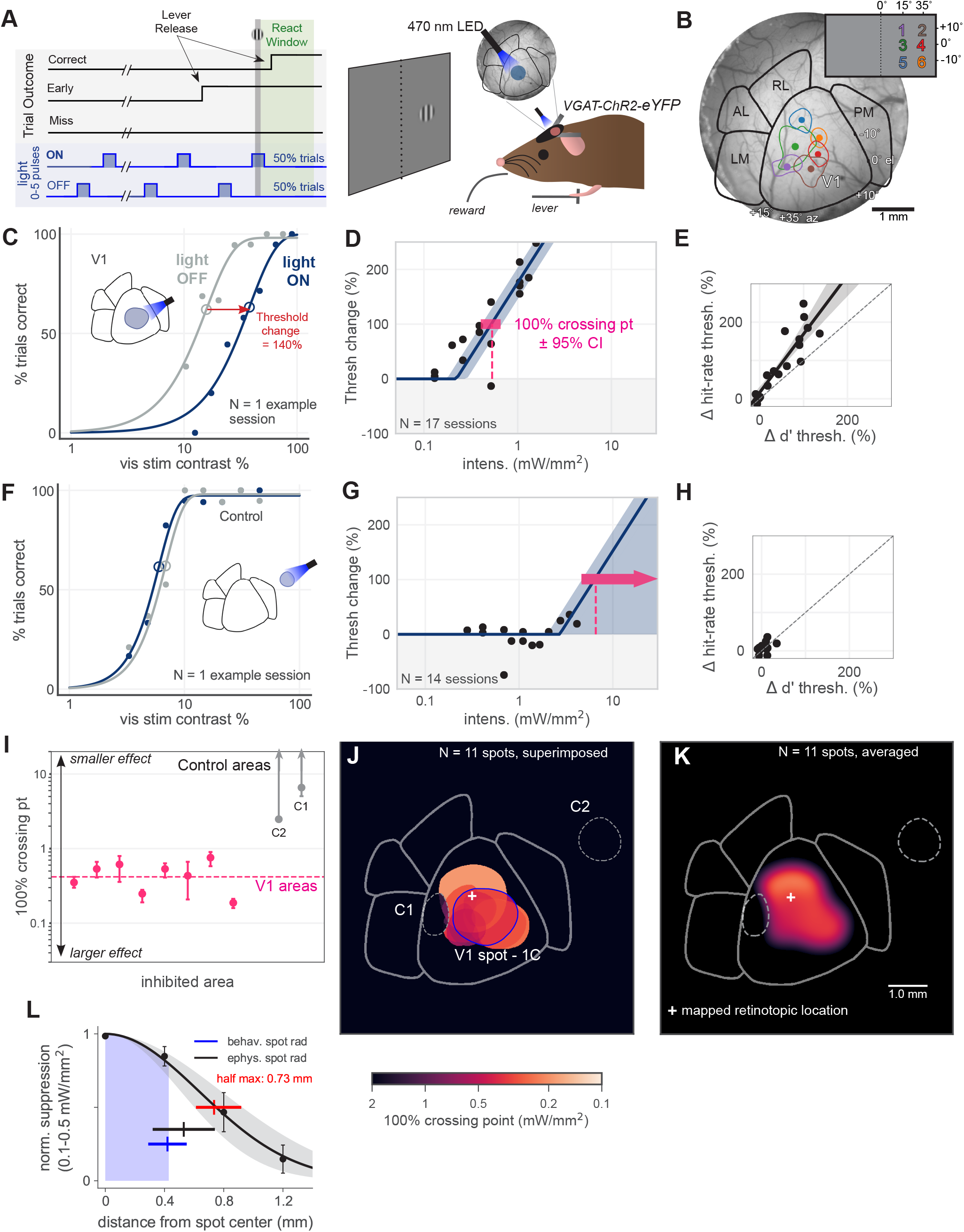
Optogenetic inhibition of V1 confirmed to affect behavior. **(A)** Schematic of the visual detection behavior task. Animals’ task was to release a lever when they detected a change in a visual stimulus. Stimuli were Gabor patches which increased in contrast, appearing on a neutral gray screen. Cortical activity was suppressed via optogenetic activation of inhibitory neurons (*VGAT-ChR2-EYFP* mouse line). On each trial, a train of light pulses was applied to the cortex (Methods). Visual stimuli were delivered either during a light pulse (light ON trials) or when the light pulse was off (light OFF trials). **(B)** Cortical areas were identified using hemodynamic intrinsic imaging of cortical responses to a set of 6 visual stimuli in different positions in visual space (Methods, ***Figure 1–Figure Supplement 1***) **(C)** LED stimulation in V1 decreases animal’s performance (single session; rightward shift indicates decreased performance). Filled circles: performance at one contrast. Open circles: threshold (62% point of fit curve). **(D)** Piecewise-linear function fit to all sessions from the spot in panel C, measuring the intensity required to double the psychometric threshold (100% crossing point, pink; mean 0.53, bootstrap 95% CI 0.40-0.66 mW/mm^2^). Slopes were fit by pooling across stimulation locations and experimental days; fitting slope separately for each day produced similar results (***Figure 1–Figure Supplement 2***). **(E)** Change in *d*′threshold increase vs change in hit-rate threshold increase for the spot in panel C (Methods, ***Figure 4–Figure Supplement 1***). **(F)** Single day example behavior from LED stimulation in a control area outside of V1, same conventions as panel C. **(G)** Same as panel D, fit to sessions from control spot in panel F **(H)** Same as panel E, fit to sessions from the control spot in panel F; mean 100% crossing point 6.6, bootstrap 95% CI 4.6-8.1 mW/mm^2^. **(I)** 100% crossing points for all V1 spots and two control spots, as determined by regression fits in E & G. Ordering of spots is arbitrary; points are offset along x axis to allow display of confidence intervals for each point. **(J)** Contours of all V1 spots and controls, color weighted by 100% crossing point. **(K)** Heatmap of effect size in V1, generated by averaging the 100% crossing point at each pixel. **(L)** Spatial fall-off of inhibition. Y-axis: suppression of visual responses measured with electrophysiology (silicon probe, four shanks), light intensity between 0.1-0.5 mW/mm^2^ (N = 4 experiments, black line: Gaussian fit, gray: 95% CI via bootstrap; data errorbars: SEM, N = 45 single units). Full-width at half max of Gaussian fit = 0.73 mm, bootstrap 95% CI 0.61-0.92 (red line). Electrophysiology spot radius: 0.53 ± 0.21 mm (black line, mean ± std dev). Behavioral spot radius: 0.42 ± 0.13 mm (blue shaded area and line, mean ± std dev, N = 30 inhibitory light spots). Since behavioral spots are smaller on average than used for these physiology experiments, the spatial effect of inhibition during behavior is likely more restricted than shown by these suppression data (black curve, red bar).

Once mice were trained to threshold, we introduced a pulsed blue LED train (470 nm; on for 200 ms, off for 1000 ms) throughout each trial. On 50% of trials (ON trials), the LED pulse occurred with the visual stimulus. (Due to latency between visual stimulus onset and first spikes in the cortex, we turned the light pulse on 55 ms after visual stimulus onset; 8-12 ms before the visual response occurred in V1 ***Figure 5–Figure Supplement 1***.) On the remaining trials (OFF trials), the visual stimulus appeared while the light was off (***Figure 1A***, 820 ms after the last light pulse was turned off).

To limit the information the animals could gain about correct response (*i.e*. that they could gain about visual stimulus onset time) from the optogenetic light pulses, the light pulse train was present on each trial, and the pulse train phase was randomized by varying the first pulse onset time after the trial began (Methods). On each trial, the visual stimulus onset time was chosen randomly (according to a geometric distribution), the light pulse train phase offset was chosen randomly (uniform distribution), and then the visual stimulus onset time was adjusted downward to be during the previous light pulse (ON trials) or after the previous light pulse offset (OFF trials). Thus, each trial, whether ON or OFF, contained a similar pulse train, and we randomly intermixed ON and OFF trials. These factors made it difficult for animals to use information about the light pulses to identify the type of trial and change their strategy (such as their perceptual criterion) from trial to trial. While some small amount of information about the visual stimulus time might in theory be available, because the visual stimulus was locked to discrete times relative to pulse onset or offset (Methods), we observed no evidence animals used light pulse information to guide responses. We found (details below): (1) the optogenetic stimulation systematically decreased, not increased, animals’ performance, (2) control-site stimulation did not change performance, and (3) sensitivity (*d*′) analyses yielded qualitatively the same results as hit-rate measures. Therefore, this behavioral change detection paradigm allows us to examine effects on visual perception due to inhibition of different mouse visual areas. Finally, while cognitive factors like attention and motivation can change over experimental sessions from one day to the next, many such factors should act similarly on both trial types. Therefore, to control for changes in attention, motivation, or perceptual criterion choice (***Macmillan and Creelman, 2004***) from day to day, we made our comparisons always between ON and OFF trials within each daily experimental session (e.g. gray vs blue curves, ***Figure 1C,F***).

### V1 inhibition confirmed to impair behavior

We first sought to validate our methods in primary visual cortex. Based on previous work (***Glick-feld et al., 2013***), we expected that inhibition of V1 during a contrast-increment change detection task would impair the animals’ behavior. Indeed, we found that inhibiting V1 during behavior, via a retinotopically-aligned spot of light over V1, produced an intensity-dependent impairment in behavior (increase in psychometric threshold; ***Figure 1C***). In order to obtain a measure of how effective the optogenetic inhibition was at impairing behavior, for each stimulation light spot we fit a piecewise-linear function to the threshold increases as a function of optogenetic light intensity. This yielded an estimate of the intensity required to produce a 100% increase in threshold (yielded the 100% crossing point; ***Figure 1D, Figure 1–Figure Supplement 2***), and allowed us to compare the behavioral impact of inhibition across light spots ***Figure 1I-K***. False alarm (early response) rates were only changed a small amount by stimulation (***Figure 4–Figure Supplement 1***, for this V1 spot: mean ± s.d. early rate increase (OFF - ON): +7.4% ± 3.7%; overall in V1: +7.7 ± 4.7%.) Note that false alarm rates vary between light ON and OFF trials, but cannot vary across contrasts, because by definition false alarms must occur before stimulus presentation — before any stimulus is accessible to perception (Methods). Since false alarm rates changed only a small amount, we expected that effects measured with sensitivity (*d*′) would be similar to those measured with hit rates. To test this, we also computed *d*′thresholds in an analogous manner to hit rate thresholds. We fit a Naka-Rushton function to *d*′values at each contrast level (Methods, ***Herrmann et al., 2012***), and did so for light ON and light OFF trials to find two thresholds for *d*′. Sensitivity threshold changes were strongly correlated with hit rate threshold changes (***Figure 1E***). We also show below that the pattern of effects with inhibition of different visual areas are similar for hit rate and for sensitivity. Because early rate changes were not large, and thresholds measured with sensitivity and with hit rate were related (for all spots in V1, Wilcoxon signed-rank test *W* = 6.0, N_1_ = N_2_ = 8, *p* = 0.09), here and below we primarily calculate threshold changes on hit rate.

Inhibiting control areas, away from visual areas, would be expected to produce little effect in this task. Indeed, inhibiting control areas in the same manner as we inhibited V1 produced no effect on threshold (***Figure 1F***). While the light OFF threshold in panel F varies from the light OFF threshold in panel C, this was not a major effect across our data (light OFF threshold for all V1 sites: mean 12.5%, 95% CI 11.3-13.6%; for all control sites, mean 9.8%, 95% CI 8.7-11%) and there was no systematic relationship between threshold change and light OFF threshold (across all V1 sites, slope of linear regression *t* = −1.2, N = 8 light spots, *p* = 0.23). To determine the minimum intensity that would be necessary to produce a 100% increase in threshold (***Figure 1G***), we fit the same piecewise-linear function (***Figure 1–Figure Supplement 2*** for slope calculations) to this data across all collected intensities. For these control light spots, the data put a lower bound on the intensity required to produce a behavioral effect. In contrast, the upper confidence interval for these plots extends upward indefinitely, as we found no intensity high enough for these light spots to cause strong behavioral effects and thus no upper bound. As with V1 inhibition, changes in false alarm rates were low (summarized in ***Figure 4–Figure Supplement 1***), and thus, with little hit rate or false alarm rate change, sensitivity (*d*′) changes were also low (***Figure 1H***).

To measure the spatial falloff of optogenetic inhibition, we used electrophysiological recording combined with optogenetic stimulation (***Figure 1L***). We measured suppression at powers used to produce behavioral effects; around the 100% crossing points seen in V1 (0.1-0.5 mW/mm^2^; N = 4 recording sessions.) The half-max radius of suppression (0.73 mm, 95% CI 0.61-0.92) was slightly larger than the half-max radius of the light spots used in the recordings (0.53 ± 0.21 mm, mean ± std dev). Thus, the light spots we used produce localized suppression around the location where light was delivered.

In these electrophysiological measurements, (***Figure 1L***) we characterized the spatial falloff of suppression of visual responses, because cortical baseline (spontaneous) firing rates are more sensitive to optogenetic manipulation than are evoked visual firing rate changes (***Glickfeld et al., 2013***; ***Histed, 2018***). We positioned an array of four electrodes so three electrodes were in V1 and the last electrode shank was in PM, which was targeted with optogenetic light. In this configuration, the farthest electrode (1.2 mm), which shows the smallest suppression, is at the center of the V1 retinotopic representation for the stimulus. As discussed below, there is little feedback effect on V1 of stimulating PM (and, in fact, at this light intensity, little feedback effect of stimulating any secondary visual area.) Recordings spanning from LM to V1 (instead of from PM to V1, as here, ***Figure 1L***) show similar spatial falloff of suppression (***Figure 5–Figure Supplement 3***, compare 50% crossing point in ***C***, yellow.) The specificity of our optogenetic manipulations is also confirmed by behavioral measurements (above: V1; other areas below), as the pattern of effects on perceptual behavior is dependent on where the light is delivered to the cortex.

Our V1 results were consistent across animals and across V1 light locations (mean 100% crossing pt 0.46, bootstrap 95% CI 0.33-0.58 mW/mm^2^; ***Figure 1I***). In total, we tested eight spots within V1, all of which produced substantially larger effects (*i.e*. lower 100% crossing points) than inhibition of control spots (N = 4 light spots, mean control spot 100% crossing point 5.1, & bootstrap 95% CI 2.3-8.0 mW/mm^2^; Mann-Whitney *U* = 0.0, V1 N = 8 light spots, control N = 4 light spots, *p* = 0.0042 two-tailed) (***Figure 1J***). Notably, inhibition of an area (control spot 1) placed within V1 but offset from the retinotopic location of the visual stimulus produced no effect (***Figure 1J***, control spot 1). To visualize the effects of inhibiting V1 across our stimulation locations (N = 10 light spots; N = 8 V1 light spots, N = 2 control light spots), we averaged the 100% crossing points at each pixel within V1 to generate a heat map (***Figure 1K***). Because this heatmap representation is influenced by the locations of the optogenetic light spots, it is not intended to be fully quantitative, but instead to provide a visual guide to assess the spatial pattern of effects given the spot locations used (all shown in ***Figure 1K*** and below, to allow comparison between visual areas.)

Together, this data confirms that inhibiting mouse V1 negatively affects an animal’s ability to perform a simple contrast-increment change detection behavior.

### Inhibiting areas lateral to V1 (LM/AL and RL) degrades contrast change detection

We next inhibited secondary visual areas, beginning with the areas LM, AL, and RL, which are lateral to V1. Our main goal was to determine whether any of the lateral visual areas could impact behavior in this task. Therefore, from here on, we refer to LM and AL together as LM/AL, as our optogenetic light spots could not easily differentiate between the two. AL is small relative to our spots’ effects (***Figure 1L***) and further, the retinotopic representation of our stimulus is on the border between these two areas (***Garrett et al., 2014***; ***Zhuang et al., 2017***).

Inhibiting these lateral areas, LM/AL and RL, produced large increases in psychometric threshold at low intensities. This resulted in small 100% crossing points (mean 100% crossing point 0.36, bootstrap 95% CI 0.16-0.59 mW/mm^2^) (***Figure 2A,B***). Inhibition of a control spot between V1 and LM (same as control spot 1, ***Figure 1***) produced no effect on behavior, even at high intensities (***Figure 2C***). The difference between effects on behavior due to inhibition of lateral areas and control areas was large (Mann-Whitney *U* = 0.0, lateral N = 6 light spots, control N = 4 light spots, *p* = 0.0071 two-tailed; ***Figure 2E***).

**Figure 2.**
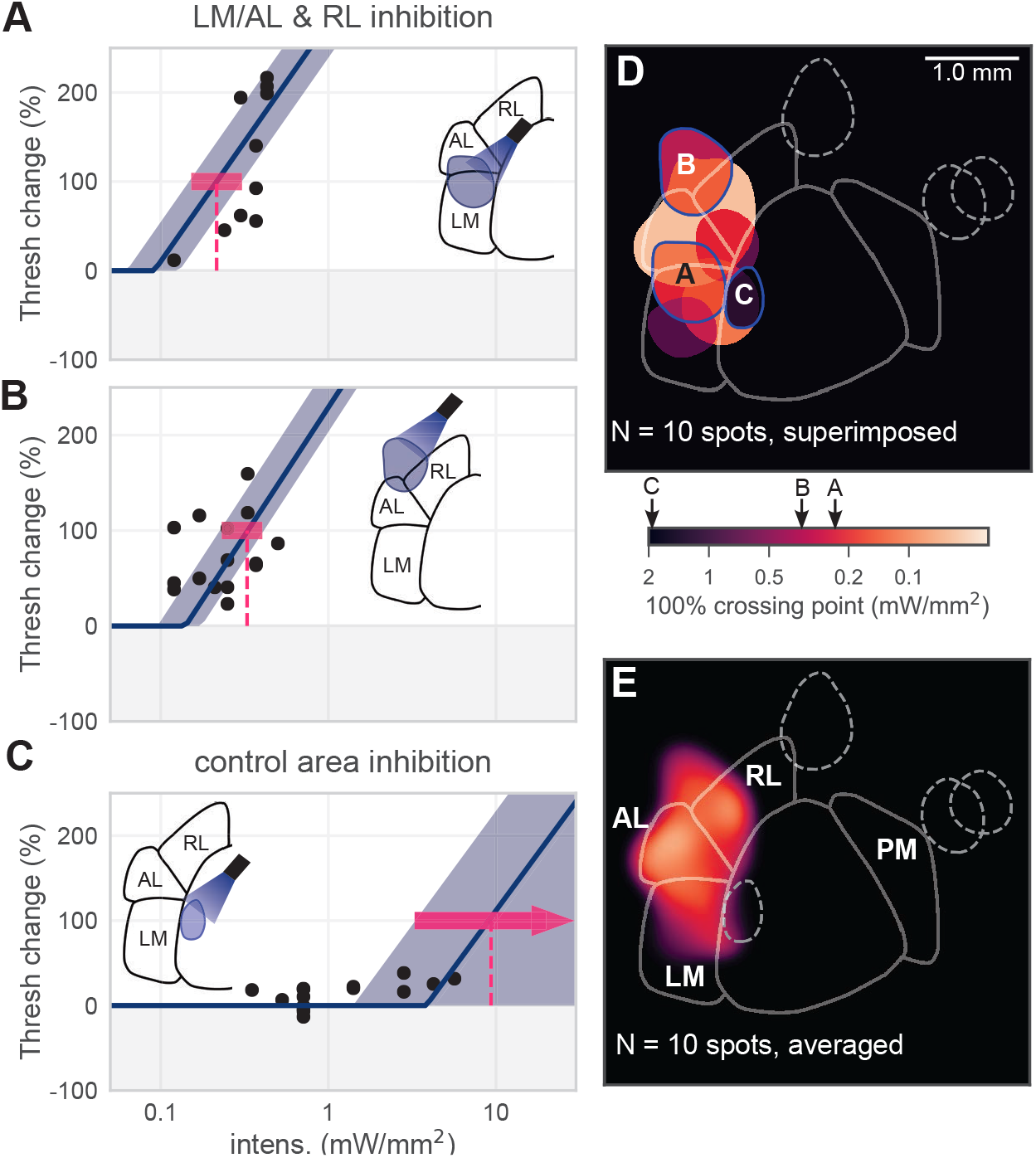
Inhibiting lateral areas degrades contrast change detection behavior. **(A)** All sessions from an example spot on the border of AL and LM, showing a large effect (mean 100% crossing pt 0.22, bootstrap 95% CI 0.15-0.31 mW/mm^2^). **(B)** All sessions from an example spot within RL, showing a large effect (mean 100% crossing pt 0.33, bootstrap 95% CI 0.23-0.40 mW/mm^2^). **(C)** All sessions from a control area between LM and V1, showing no effect even at high intensities (mean 100% crossing pt 9.3, bootstrap 95% CI 3.3-10 mW/mm^2^). **(D)** Map of spots within LM/AL/RL and control locations, colored by 100% crossing pt. Crossing points from panels A-C are shown on colorbar (black arrows). (Note slight difference for visual display in colorbar extents compared to ***Figure 1J-K***). **(E)** Heatmap of effect size, generated by averaging the 100% crossing point at each pixel; pixels with no data are colored black.

The piecewise-linear function we used fixed the slope of each curve. While there was some variability in slope across data points (***Figure 2A,B***), we used this approach because in V1 allowing slopes to vary in the fits did not appreciably change the estimated crossing points (***Figure 1–Figure Supplement 2***).

In sum, inhibition of these secondary visual areas, lateral to primary visual cortex, resulted in substantial degradation of the animals’ ability to perform the contrast change detection task.

### Inhibiting PM produces smaller effects than inhibiting lateral areas (LM/AL & RL)

In addition to inhibiting areas within LM, AL, and RL, we also inhibited locations medial to V1, within area PM. We found that on average, inhibiting spots within PM produced little effect on behavior (mean PM 100% crossing point 1.7, bootstrap 95% CI 0.67-2.7 mW/mm^2^) (***Figure 3A, Figure 3B***). Inhibiting control areas located outside PM had small or undetectable effects on behavior, even at relatively high light intensities (≥5 mW/mm^2^, ***Figure 3C***). Of five light spots targeted to PM, two showed an obvious effect, and the effect was of medium size compared to V1 and the lateral areas (***Figure 3D,E***).

**Figure 3.**
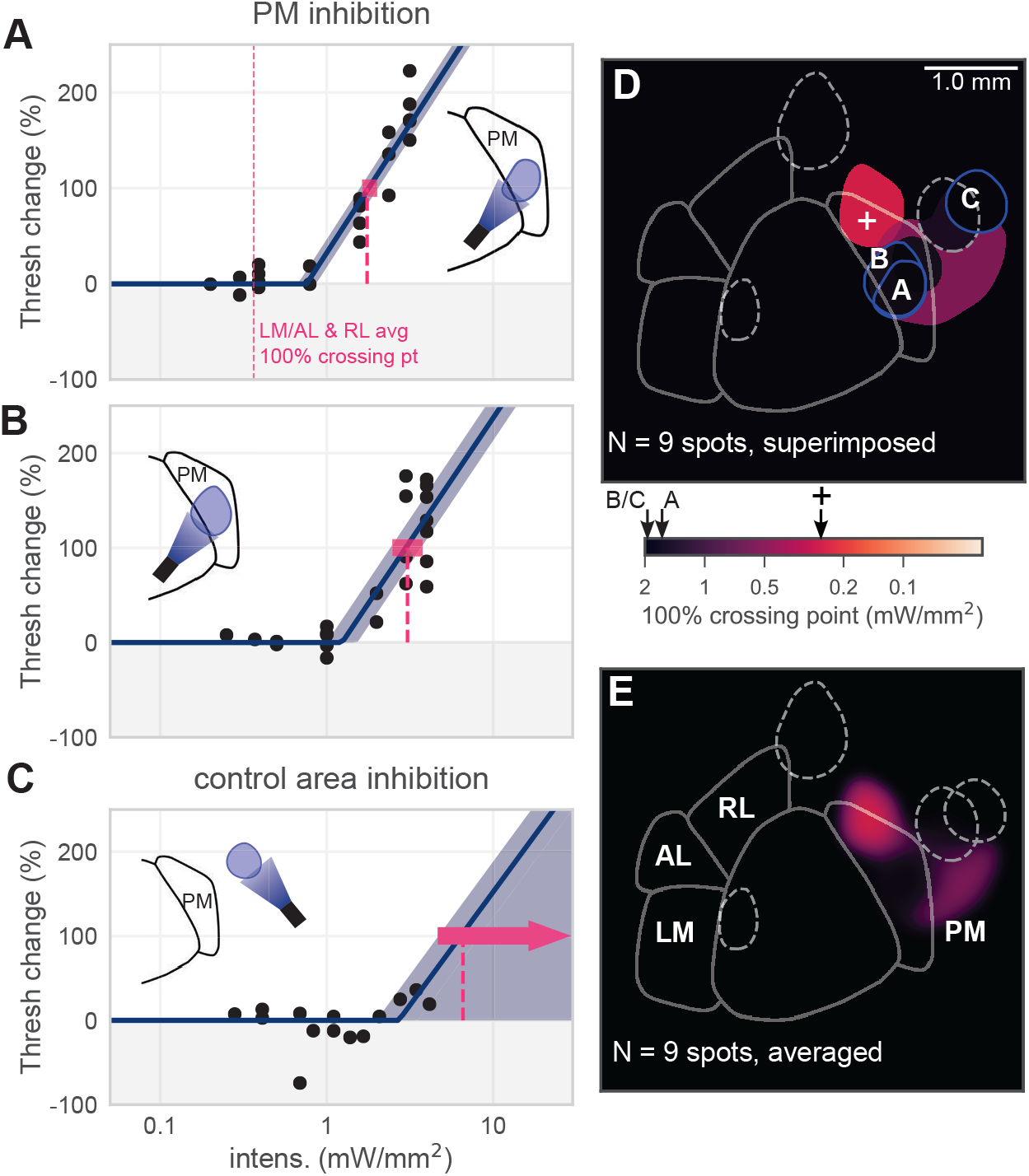
Inhibiting medial areas produces weaker effects on behavior. **(A)** All sessions from an example spot within PM, which produced a weak effect (mean 100% crossing pt 1.7, bootstrap 95% CI 1.6-2.0 mW/mm^2^). Vertical pink dashed line represents average crossing point from LM/AL and RL (0.36 mW/mm^2^). **(B)** All sessions from an example spot a second spot within PM, which also produced a weak effect (mean 100% crossing pt 3.1, bootstrap 95% CI 2.5-3.5 mW/mm^2^). **(C)** All sessions from a control area outside PM, showing very little effect even at high intensities (mean 100% crossing pt 6.6, bootstrap 95% CI 4.6-8.1 mW/mm^2^). **(D)** Map of spots within PM and control locations, colored by 100% crossing pt. Crossing points from panels A-C are shown on colorbar (black arrows). **(E)** Heatmap of effect size, generated by averaging the 100% crossing point at each pixel.

Most PM spots produced small effects on behavior, with the one PM spot that produced a moderate effect at the most anterior point of PM. That area of PM has a retinotopic representation that is slightly more temporal in azimuth (***Garrett et al., 2014***; ***Zhuang et al., 2017***) than the stimulus we used (25°azimuth). To determine if smaller PM effects relative to LM/AL/RL were due to the retinotopic position of the stimulus, we measured effects of inhibiting anterior PM while animals performed the same task with stimuli at 45-65°azimuth. (***Figure 3–Figure Supplement 1***). We found small or nonexistent effects for these stimuli. Across 6 PM light spots, using both 25° and 45-65° azimuth stimuli (***Figure 3–Figure Supplement 1***), despite two light spots that showed a moderate effect, the mean PM effect was small. In sum, for the same stimulus used for V1 and LM measurements, and for a more eccentric stimulus that may be better represented in PM, effects of inhibiting PM were significantly smaller than those that arose from inhibiting LM/AL and RL.

### Inhibition of LM/AL/RL produces similar or larger effects on behavior as inhibiting V1

We compared the strength of behavioral effects of inhibition across areas by computing the mean effect (mean 100% crossing point, the measure of how much light is required to produce a fixed degradation in behavior) in each region: lateral areas, PM, and V1. We found that the mean effect was similarly strong (smaller crossing point intensity) in the lateral areas and in V1 (***Figure 4A***; Mann-Whitney *U* = 17.5, lateral N = 6 spots, V1 N = 8 spots, *p* = 0.22 two-tailed), while the mean effect across all PM spots was weaker (larger crossing point intensity) than in V1. The average response heat maps (***Figure 4***; lighter colors are stronger effects on behavior and thus smaller crossing point intensities) show the spatial extent of these effects given the spot positions we tested.

**Figure 4.**
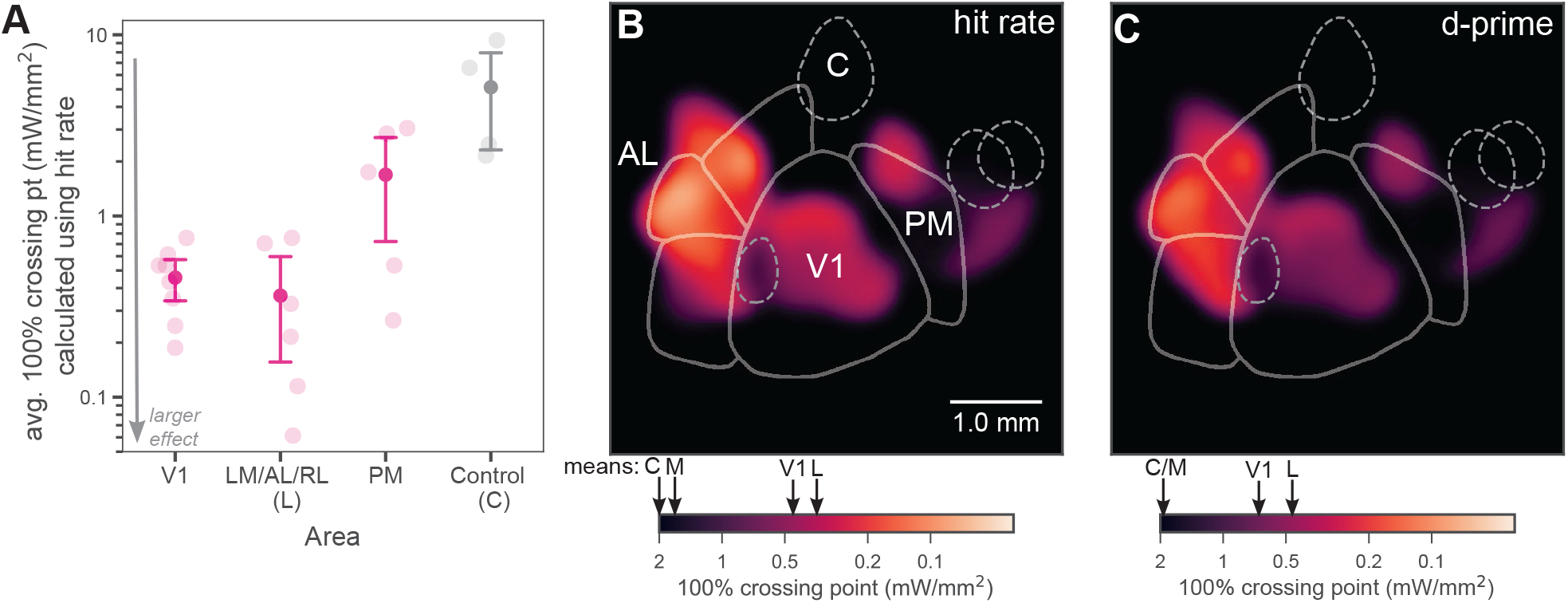
Inhibiting lateral areas produces larger effects than inhibiting PM. **(A)** Average 100% crossing point (± bootstrap 95% confidence interval) across all spots within a given area (V1 / Lateral / Medial / Control). V1 and lateral areas (LM/AL/RL) require very low light intensities to double threshold (mean 0.46, bootstrap 95% CI 0.33-0.58 mW/mm^2^ and 0.36, CI 0.16-0.59 mW/mm^2^, respectively). Inhibited areas within PM required much higher intensities (mean 1.7, bootstrap 95% CI 0.67-2.7 mW/mm^2^) to achieve the same effect, and control areas required very high intensities (mean 5.1, bootstrap 95% CI 2.3-8.0 mW/mm^2^). **(B)** Heatmap of all inhibited spots, colored by weighted hit-rate 100% crossing point. Light intensity required to produce effects in lateral secondary areas is similar to or larger than that needed in V1. Mean crossing points from spots within V1, LM/AL & RL (L), PM (M) and control (C) are represented on colorbar with black arrows. **(C)** Heatmap of all inhibited spots, colored by weighted *d*′100% crossing point. When 100% crossing point is calculated in terms of *d*′, spots in V1 and lateral areas still require much less intensity to double threshold (means 0.68 and 0.48 mW/mm^2^, respectively) than spots in PM and control areas (means 3.0 and 6.1 mW/mm^2^, respectively). Mean crossing points are represented on the colorbar using the same notation as ***B***.

Perceptual behavioral performance can be affected both by changes in sensitivity, the amount of information subjects can use about the stimulus change, or by changes in perceptual criterion, the subject’s willingness to respond “yes” (in this case, willingness to release the lever). A more permissive criterion increases hit rate at the cost of a higher false alarm rate. In the V1 data (***Figure 1***), we found that optogenetic inhibition caused little change in false alarm rate, and thus sensitivity (*d*′) changes mirrored the hit rate changes. We asked whether there were similar or different changes in false alarm rates between inhibiting V1 and secondary visual areas. We found no significant difference in early rates between areas (mean ± s.d. early rate increase (OFF - ON) V1: 7.4% ± 4.7%; LM/AL & RL: 9.1% ± 6.0%; PM: 6.6% ± 5.0%; control: 6.7% ± 5.2%) (***Figure 4–Figure Supplement 1***). Given the small effect on early rate, which also varied little between areas, we expected little change in the pattern of our results when measured in terms of sensitivity. Indeed, calculating 100% crossing points in terms of sensitivity (*d*′) had no effect on the observed differences between areas (V1: Wilcoxon signed-rank test *W* = 6.0, N_1_ = N_2_ = 8, *p* = 0.09; LM/AL and RL: Wilcoxon signed-rank test *W* = 5.0, N_1_ = N_2_ = 6, *p* = 0.5; PM: Wilcoxon signed-rank test *W* = 6.0, N_1_ = N_2_ = 5, *p* = 0.66; Control: Wilcoxon signed-rank test *W* = 4.0, N_1_ = N_2_ = 4, *p* = 0.72) (***Figure 4C, Figure 4–Figure Supplement 2***).

We next examined changes in animals’ response bias, or willingness to respond. There are several ways to characterize response bias, and we examined two different measures: absolute criterion and relative criterion ***Macmillan and Creelman*** (***2004***). Absolute criterion (*c*) is the distance of the decision threshold, in the stimulus space of signal detection theory, from the idealized zero point in that space. Absolute criterion varies with both hit rate and false alarm rate. When, as in our data, false alarm rate change is small compared to change in hit rate, changes in absolute criterion (*c*) are primarily due to changes in hit rate (*c* = (*z*(*H*) + *z*(*FA*))/2, where *H* and *FA* are the hit and false alarm rates, and *z*(·) is the inverse normal CDF). As expected, absolute criterion *c* was changed by optogenetic suppression (data pooled across areas, regression of hit rate change vs absolute criterion change, N = 300 sessions, *t* = −22, *p <* 0.001). However, for this task, relative criterion (c’; = *c*/*d*′) is a more appropriate measure of bias, as relative criterion accounts for changes in sensitivity. Indeed, we saw little change in relative criterion with optogenetic inhibition (c’ change, laser on - laser off: median 0.016, inter-quartile range 1.26, not sig. different from zero: Wilcoxon W = 3.3 × 10^4^, *p* = 0.6; linear regression of c’ change on hit rate change *t* = −0.87, N = 300 sessions, *p* = 0.33). Finally, the false alarm rate itself is another way to measure response bias in a temporally extended change detection task like this, since false alarm rate reflects subjects’ willingness to respond for a single stimulus, the fixed visual stimulus (of zero contrast) before the contrast increment. We found above that the optogenetic perturbation only slightly affected false alarm rates. Thus, the systematic effects of optogenetic suppression on behavior are primarily due to decreases in sensitivity, not response bias; cortical suppression reduced animals’ ability to detect the presence of the stimulus.

Effects of optogenetic inhibition could affect both sensory responses or, in principle, animals’ motor responses. One way to determine whether our optogenetic inhibition affected motor responses is to vary the motor response the animal uses to report a stimulus change, while keeping the sensory stimulus the same. We did this by training an animal to perform the task using the forepaw ipsilateral to the visual stimulus (in contrast to all other behavioral data in this work, where animals were trained to use their contralateral forepaw). If inhibition affected motor responses, we would expect to see a different pattern of behavior effects when animals’ motor response type was changed. Instead, we saw similar effects in V1, LM/AL, and RL regardless of whether the ipsilateral or contralateral paw was used to report change (***Figure 4–Figure Supplement 3***). While these data cannot rule out some motor contribution, they are evidence that inhibition of these visual areas produced an effect primarily on sensory information, not motor execution.

Finally, the visual stimulus location we used for the experiments above (+25° azimuth, 0° elevation) is near the border of area PM, and it might be possible that a stimulus in a more temporal location (*i.e*. displaced in the horizontal direction away from the animal’s nose) could produce a larger neural response and perhaps reveal larger effects of inhibition. However, when we measured the effect of inhibition with a visual stimulus at a range of more temporal azimuths (+45-65° azimuth, 0° elevation), we found similarly small effects on behavior as with the more central stimulus (***Figure 3–Figure Supplement 1***). Thus, even across a range of stimulus retinotopic positions, the effect of PM inhibition on this perceptual behavior is not large.

In sum, inhibition of areas lateral to V1 (LM/AL and RL) produces the strongest effects on contrast-increment change detection, as large or larger than in V1. The fact that some secondary visual areas (the lateral areas) show these strong effects implies that V1 is not being directly decoded by remote areas, but instead that information must pass through these secondary visual areas before being used for contrast change detection.

### Behavioral changes are primarily due to direct suppression, not feedback

Although we found above that inhibiting secondary visual areas impacted animals’ contrast-detection behavior, it could have been possible that the effect of inhibiting secondary visual areas was merely to suppress V1 activity, in a similar way to inhibiting V1 itself. To explore this possibility, we performed a series of electrophysiology experiments, where we inhibited in V1, lateral areas, or medial areas while simultaneously recording at the location where optogenetic light was delivered (Methods; ***Figure 5A***).

**Figure 5.**
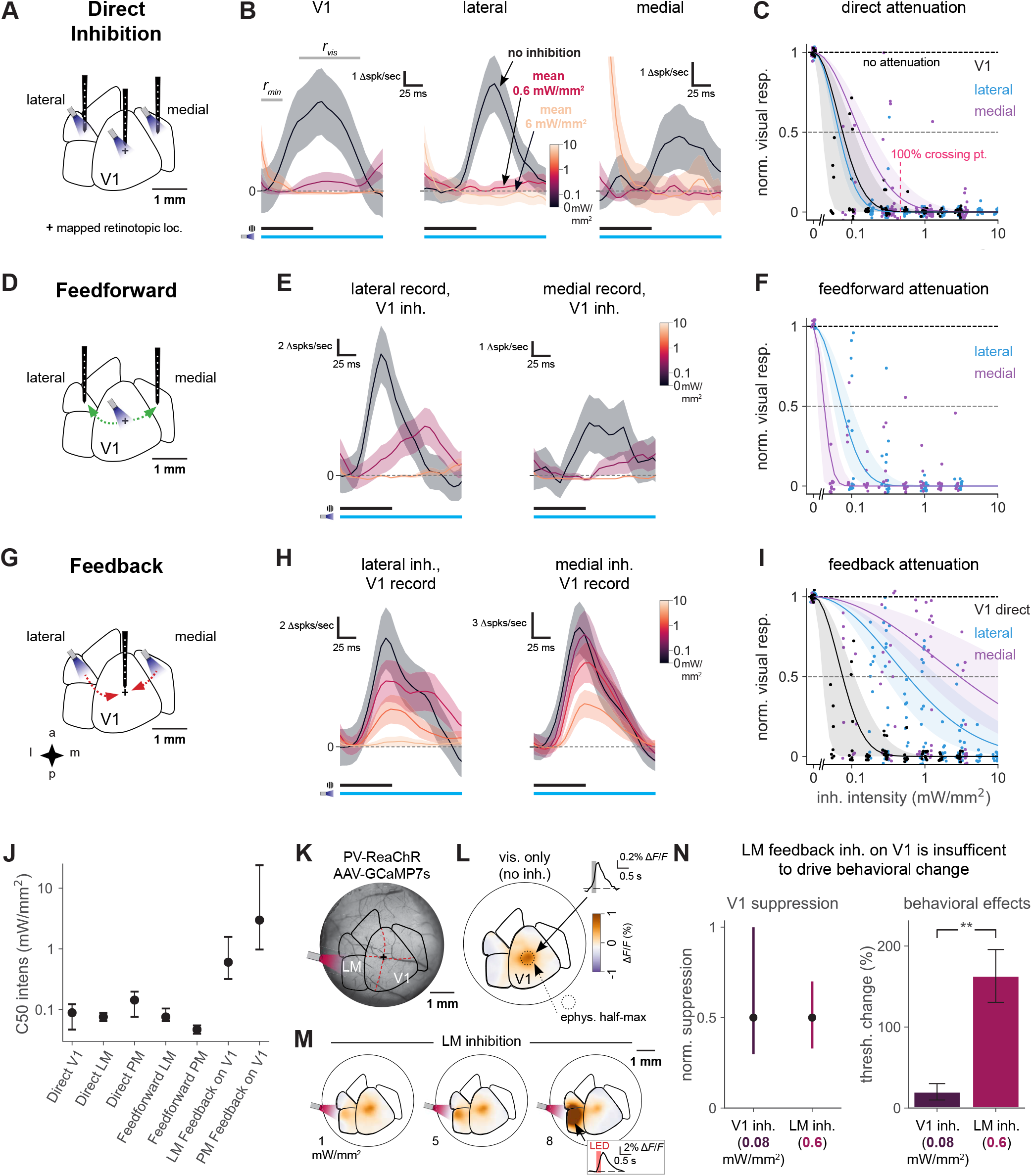
Direct activity reduction in secondary areas, not exclusively feedback suppression onto V1, accounts for degradation of behavioral performance. **(A)** Schematic of direct optogenetic inhibition of V1, lateral, and medial areas. Plus sign: visual stimulus at a mapped retinotopic location in V1. **(B)** Average visual responses in each cortical area with increasing amounts of optogenetic inhibition (0, 0.6, and 6.0 mW/mm^2^). Responses are subtracted by baseline r_*min*_, a 25 ms period where optogenetic inhibition has reduced spontaneous neural activity but before visually evoked spikes arrive to the cortex. r_*vis*_: visual response period for analysis in C. Black and blue bars: duration of visual stimulus and optogenetic inhibitory stimulus, respectively. Vertical scale bars in panels vary, to more clearly illustrate relative suppression with optogenetic stimulation; quantification in panel C. The rightmost panel shows at the highest power a transient associated with ISN dynamics (***Sanzeni et al., 2020, Figure 5–Figure Supplement 2***), which has ended by the time the visual response analysis period begins. V1, lateral, medial panels: N = 14, 6, 7 single units. **(C)** Summary of direct inhibition effects on visually evoked responses for all intensities. One point is shown for every unit and every intensity level (4-5 intensities per unit). Normalized visual response of 1 is no attenuation (black dotted line). Points are jittered slightly in both x and y directions for visual display, including at zero intensity where *y* = 1 for all points. Pink dotted line: Mean LED intensity for V1 100% behavior threshold change. Solid curves: gaussian fits, shaded region: 95% CI via bootstrap. 50% suppression level shown by lighter dashed line. **(D)** Schematic of inhibition of feedforward V1 connections to secondary visual areas (green arrows). **(E)** Average visual responses measured in lateral areas and PM with increasing optogenetic inhibition applied to V1; conventions as in panel B. Lateral, medial panels: N = 4, 6 single units. **(F)** Summary of feedforward effects. Conventions as in panel C. **(G)** Schematic of feedback suppression of V1 during inhibition of secondary visual areas (red arrows). **(H)** Average visual responses measured in V1 with increasing optogenetic inhibition applied to lateral areas or PM. Lateral, medial panels: N = 26, 16 single units. **(I)** Summary of direct V1 inhibition versus feedback suppression for all intensities tested. **(J)** Intensities (of inhibitory optogenetic stimulus) that generate 50% visual response suppression for all methods of inhibition (direct, feedforward, and feedback) in all areas (V1, lateral areas, and PM). Errorbars: 95% CI; all taken from intersection points with colored lines, shaded regions in panels C,F,I. Higher intensities are required to produce suppression in V1 through feedback than through either direct or feedforward suppression of V1 to other areas. **(K)** Schematic of GCaMP7s imaging with LM inhibition in a mouse expressing ReaChR in all PV cells (PV-Cre;floxed-ReaChR mouse). **(L)** GCaMP7s response to flashed Gabor visual stimulus. Fluorescence map image is calculated by taking the frame-by-frame difference, approximating a spike-deconvolution filter; dF/F response without differencing is shown in inset. V1 activation is restricted to the retinotopic location of the stimulus. **(M)** Responses to the visual stimulus paired with LM inhibition at 3 intensities. Increases in activity (orange) in LM/AL likely reflect increased firing of inhibitory neurons expressing ReaChR (inset: LM/AL light response timecourse has a decay consistent with GCaMP7s offset dynamics). **(N)** Intensity of optogenetic stimulation required to suppress V1 activity directly is much less than needed to achieve same suppression by illuminating LM. Intensites in both areas were chosen to produce the same mean suppression. Direct: 0.08 mW/mm^2^, suppression 0.5, CI 0.3-0.9, N = 4 recording sessions. Feedback: 0.6 mW/mm^2^, suppression 0.5, CI 0.4-0.7, N = 6 recording sessions). Behavioral effects at these powers are very different, indicating that behavioral effects of LM suppression arise principally by changing LM responses, not by feedback inhibition of V1. (V1 inh. at 0.08 mW/mm^2^, N = 8 light spots; lat. inh. at 0.6 mW/mm^2^, N = 6 spots; Mann-Whitney *U* = 1.0, one-sided, *p <* 0.002).

We first examined how visual responses were suppressed at the site where optogenetic light was delivered. We found that optogenetic stimulation of inhibitory neurons reduced or abolished visual responses at the site of light stimulation (units recorded from V1, AL/LM border or in PM; ***Figure 5B***). We found that LED intensities that produced behavioral effects (in V1, 0.46 mW/mm^2^ to produce a 100% threshold change) completely suppressed neural activity directly under the LED (V1, bootstrap 95% CI 96-100% suppression of visual response without light, ***Figure 5C***).

We next examined visual responses to contrast increments in the different visual cortical areas. Peak visual responses were similar in size between V1 and lateral areas, and both were significantly larger than peak responses in PM (***Figure 5–Figure Supplement 1A,B***). We measured visual stimulus latency (time to half-max response, sigmoidal fit between r_*min*_ to r_*vis*_) in V1 to be 65 ms (bootstrap 95% CI 63-67 ms). There was a small delay in response in the lateral areas relative to V1 (time to half max response = 73 ms, bootstrap 95% CI 70-77 ms) but a notably larger delay for PM (time to half-max response = 90 ms, bootstrap 95% CI 87-93 ms, ***Figure 5–Figure Supplement 1C***).

Since mouse V1 operates as an inhibition stabilized network (***Sanzeni et al., 2020***), at lower light intensities many inhibitory neurons should decrease their firing while excitatory cells also decrease their firing. However, at high light intensities some inhibitory neurons (especially the narrow-waveform inhibitory cells, ***Sanzeni et al., 2020***) increase their firing. To examine differences in baseline firing rate change due to stimulation, we sorted units by waveform width (***Figure 5–Figure Supplement 2A,B***), finding a bimodal distribution of spike waveforms (***Figure 5–Figure Supplement 2C***), as in ***Sanzeni et al***. (***2020***). For simplicity in the analyses below, we dropped narrow-waveform units (likely the narrow-spiking basket cells ***Speed et al., 2019***; ***Sanzeni et al., 2020***), thereby eliminating units that increased their baseline rates to inhibitory stimulation. For analysis of visual responses below, we focused on wide-waveform units (excitatory and also likely some non-basket inhibitory cells). Finally, we saw that secondary areas — lateral areas and PM — displayed signatures of inhibition stabilized networks, supporting the idea that secondary visual cortical areas also operate in the inhibition-stabilized regime (***Figure 5–Figure Supplement 2D-L***).

As V1 is a direct recipient of visual input to the cortex, and sends projections to other visual areas, we expected that suppressing V1 would reduce responses in other visual areas by blocking feedforward flow of visual information. Indeed, inhibiting V1 while recording downstream visual responses in either lateral areas or PM (***Figure 5D***) showed that inhibiting V1 strongly decreased visual responses in both secondary areas (***Figure 5E***). We saw this effect at LED intensities comparable to those that produced 100% changes in behavioral threshold, and the feedforward suppression was even stronger at higher intensities (***Figure 5F***). These measurements support the idea that the majority of the visual information seen in secondary visual areas passes through V1.

We next examined whether our behavioral effects in secondary areas originated solely from local suppression of visual responses or if they could have arisen from feedback suppression of V1. That is, we studied whether inhibition of secondary visual areas produced behavioral deficits due to suppression of responses in the secondary area itself, or whether inhibition could have potentially produced behavioral deficits merely by acting on the V1 responses via feedback connections. To study this, we recorded in V1 and simultaneously inhibited lateral areas or PM while awake animals were passively viewing the visual stimulus used for behavior (***Figure 5G***). We saw that optogenetic suppression of lateral areas did partially reduce V1 responses (***Figure 5H, left***). At light intensities that produced behavioral effects (0.36 mW/mm^2^), inhibiting LM/AL produced moderate (39%, 95% CI: 24-57%) suppression in V1. However, the same light stimulation nearly eliminated visual responses in LM/AL (i.e. 100% suppression). Thus, the principal effect of stimulation of LM/AL was to suppress LM/AL, with a smaller effect of suppression in V1 (***Figure 5C,I,J***).

In contrast to the feedback effects on LM/AL, we found that suppression of medial areas had little to no feedback effect on V1 visual responses, even at the highest intensities we used (***Figure 5H, right***).

To characterize the spatial extent of feedback effects on V1, we used widefield calcium imaging. We expressed viral GCaMP7s in all neurons via multiple injections into a transgenic mouse expressing ReaChR-mCitrine in all PV interneurons (i.e PV-Cre;ReaChR-mCitrine mouse; AAV-GCaMP7s injections; average green fluorescence due to GCaMP shown in ***Figure 5K***). We imaged GCaMP responses with and without LM inhibition while the animal was shown the flashed Gabor stimulus (***Figure 5K***). As expected, GCaMP responses were observed at the V1 retinotopic location corresponding to the stimulus (***Figure 5L***. When the visual stimulus was paired with LM inhibition, we saw little change in the spatial extent of V1 response and slight reduction in the amplitude of the GCaMP response (***Figure 5M***), consistent with the feedback suppression observed in physiology when LM was stimulated at high powers. At the site of the optogenetic light, we saw increases in fluorescence, with a decay time constant consistent with a GCaMP response (***Figure 5M***, inset; compare to V1 response decay, ***L*** inset), presumably due to activation of inhibitory neurons at powers high enough to profoundly suppress excitatory cells, exit the cortical ISN regime, and produce substantial inhibitory activation (***Sanzeni et al., 2020; Li et al., 2019 and Figure 5–Figure Supplement 2***). Compared to the physiological results, in these imaging data we stimulated only PV neurons, and also these results could be influenced by the spatial location of cells infected by the GCaMP virus. Despite this, these data support the idea that the spatial pattern of V1 responses is not changed by LM suppression, and these data are consistent with the physiological measurements (***Figure 5I***).

Finally, to directly compare the effects on both areas (V1 and LM/AL) of optogenetic suppression of LM/AL, we examined V1 and LM/AL stimulation intensities that produced a matching magnitude of V1 suppression. Delivering 0.08 mW/mm^2^ to V1 and 0.6 mW/mm^2^ to LM/AL, while in both cases recording in V1, produced similar levels of suppression on the visual responses of V1 neurons (***Figure 5N, left***; these data were drawn from the Gaussian fits shown in ***Figure 5C,I***). We were then able to examine in our behavioral data how the two types of stimulation (0.08 mW/mm^2^ to V1; 0.6 mW/mm^2^ to LM/AL), which affected V1 responses similarly, affected animals’ behavior. LM/AL stimulation, and not V1 stimulation, produced large degradation of animals’ contrast change detection behavior (***Figure 5N, right***). We also compared direct and feedback V1 suppression across 0.4-0.8 mm spans of cortex, at both 0.1 and 0.3 mW/mm^2^, and found that feedback suppression of V1 neural activity caused by LM direct inhibition was always two to five times lower than direct suppression of V1 neural activity ***Figure 5–Figure Supplement 3***.

Together, these observations argue that when lateral secondary areas LM/AL are suppressed, it is not the change in V1 responses that drives the behavioral effect, since stimulating V1 itself at an intensity necessary to produce the same V1 change has no effect on behavior. Instead, it likely is the profound suppression of LM/AL activity (bootstrap 95% CI 99-100% suppression) at this light intensity that leads to the behavioral degradation. In sum, though inhibition of lateral areas does suppress activity in V1, feedback suppression alone could not explain the observed behavioral results.

## Discussion

This work examines the effect of inhibiting secondary visual areas in *VGAT-ChR2-EFYP* mice performing a contrast-increment change detection task. Neurons in V1, lateral, and medial secondary visual areas change their firing rate in response to changes in stimulus contrast, but only suppressing V1 and the lateral areas produced a large impact on animals’ performance in the task. Inhibiting areas lateral to V1 sometimes caused deficits larger than those seen when inhibiting V1 directly. Behavioral effects were due to changes in sensitivity (*d*′) without large changes in response bias. When inhibiting the secondary areas, light intensities that produced behavioral changes produced the largest changes in neural firing in the secondary areas themselves, not in V1 via feedback suppression. Taken together, our results provide evidence that while substantial information about the contrast changes is present in V1, secondary areas AL/LM and RL form an essential part of the neural circuit that controls this behavior.

### Behavioral changes are consistent with sensory, not motor representations, and are spatially specific

We performed a control experiment to determine whether inhibition affected sensory signals rather than motor planning or execution, and a second control to determine if our minimal PM effects could have been created by using a stimulus that was not well-represented in the retinotopic map in PM. First, if our inhibition affected motor performance, we would have expected that such an effect would vary based on whether animals were using the contralateral or ipsilateral forepaw. Instead, we found that training animals to perform the task with their ipsilateral forepaw produced similar results as when animals used their contralateral forepaw (***Figure 4–Figure Supplement 3***). Second, we examined the possibility that our observed PM effects were weaker because PM may contain a weaker representation of the nasal than the temporal visual field. To rule this out, we measured the effect of inhibiting PM during the same task but performed with a stimulus further into the temporal visual field (45-65°azimuth relative to midline), where previous studies have found robust retinotopic representations in PM. Using this stimulus, we found similar, or smaller, magnitude effects as with our 25°eccentricity stimulus (***Figure 3–Figure Supplement 1***, arguing that the weaker effects in PM relative to lateral areas were not due to the location of the visual stimulus.

Our results examine the effect of inhibiting the cortex at a variety of different locations, and rely on several observations to establish spatial specificity. First, our key finding is that inhibiting lateral visual areas, LM and/or AL as well as RL, produces similar-sized, or larger, effects on behavior as inhibiting V1. The cortical distance involved between the lateral areas and V1 (approximately 2 mm) and the fact that several control spots with equally distant spacing (as well as PM locations) produced little or no effect, is itself evidence that our effects are due to inhibition at the site where the light was positioned. Second, the spatial specificity observed in behavioral experiments is supported by our neurophysiological recordings. We observed that the evoked visual response showed a spatial falloff consistent with, though slightly larger than, the light intensity patterns, or spots, that we used (***Figure 1L***). Recording neural responses at several locations across V1 confirmed that the spatial extent of suppression in V1 was similar for direct V1 activation and for V1 suppression when LM/AL was inhibited, though the direct suppression was significantly stronger across V1 than feedback suppression (***Figure 5 and Figure 5–Figure Supplement 3***). Imaging V1 responses with GCaMP (***Figure 5***) also showed the V1 representation was approximately the size of the area directly suppressed by our light spots, and that AL/LM suppression affected an area of similar size. In sum, these characterizations of the spatial extent of suppressive effects confirm that LM/AL suppression produces effects on behavior independent of V1.

Another factor that may have changed our intensity thresholds, but did not affect the pattern of results across areas, is variability in effect size due to stimulation spot size. Even if there is some association of effect size with spot size, considering a subset of spots with similar sizes confirms our effects (e.g. ***Figure 2D***, compare “A”, “B”, and the control spots shown in dashed lines; also see ***Figure 3D***, regions marked “B”, “C”, etc.). In one case, we used a small spot to separate AL/LM from V1 (***Figure 2D***, spot “C”), and inhibiting there produced no discernible effect on behavior up to the highest power that we used (***Figure 2C***), though the spot area was not much smaller than other LM/AL spot areas, which produced effects at much lower light intensities (***Figure 2D***). Finally, the heatmaps we use to display the spatial extent of behavioral effects (e.g. ***Figure 4B,C***) are limited to showing effects at the locations where we sited stimulation spots. Therefore, while they cannot capture the potential effect at every pixel location, they are useful to visualize average effects across areas.

### Roles of mouse cortical visual areas in contrast perception behaviors

These findings show not only that lateral areas LM/AL/RL play a role in this behavior, but also that medial areas like PM do not play a major role. (Some visual cortex parcellations include an area, anteromedial or AM, between PM and RL, ***Garrett et al., 2014; Zhuang et al., 2017***, but our conclusions do not significantly depend on whether our medial regions are part of an anterior section of PM, or a separate area AM.) Both lateral and medial areas have previously been shown to encode different features of visual stimuli, such as spatial and temporal frequencies (***Andermann et al., 2011; Murakami et al., 2017***), and thus might play object identification or localization roles, as in the ‘what’ and ‘where’ pathways of the primate visual system (***Mishkin and Ungerleider, 1982***). In this case, both lateral and medial areas might be representing different features of the visual world, and other types of behavioral decision-making might expose the role of PM (see ***Jin and Glickfeld, 2020***). On the other hand, ***Siegle et al. (2019***) constructed a hierarchy of visual areas based on electrophysiology data (LM->RL->LP->AL->PM->AM), and predicted that medial areas (PM/AM) were higher areas within the hierarchy, which serve to amplify or decode change-related signals. Our data argue against that hypothesis. If PM/AM were important for processing signals for decisions about contrast changes, inhibiting them should also serve to impair behavior, and we found only weak mean effects.

A recent study (***Jin and Glickfeld, 2020***) lends further support to the idea that PM is not crucial for contrast-perception or change detection tasks. Jin and colleagues found, as we did, that lateral areas are involved in contrast change detection tasks, and medial area inhibition has a smaller effect. Our electrophysiological and imaging measurements go beyond their behavioral results, and show that the effect of inhibiting secondary visual areas is not caused by effects on V1 but substantially caused by changes in the visual signals in the secondary areas. Our behavioral results differ in one way from theirs: on perceptual sensitivity and criterion. While ***Jin and Glickfeld*** (***2020***) found little or no change in sensitivity due to inhibiting PM, as we did, they found an effect on false alarm rates and criterion when PM was inhibited. Because subjects have some ability to choose their own criterion in perceptual behaviors (***Macmillan and Creelman, 2004***), it seems possible that this difference in false alarm rate is an effect of behavioral training. In sum, the lack of sensitivity change when PM is suppressed argues that PM is not part of the circuit that controls behavior in this contrast perception task, while V1, LM/AL and RL do play important roles in this behavior.

### Brain area interactions during perceptual behaviors

Our results shed light on the inter-area circuits used in a perceptual behavioral task. Recent work has shown that mouse premotor cortex is involved in downstream processing of sensory signals (***Wu et al., 2020; Zatka-Haas et al., 2020***), but it has been unclear whether secondary visual areas are involved in all or most sensory tasks that involve V1. Our data argue that lateral secondary areas are not bypassed during behavior, and therefore suggests the secondary areas’ activity is decoded into a choice or decision representation via those areas’ projections to downstream regions. Evidence for choice or decision signals has been found in areas including frontal and premotor cortex and the striatum, but not in either V1 or secondary areas (***Steinmetz et al., 2019***), arguing that the decoding happens outside both V1 and the secondary areas. However, there are several caveats to this sequential interpretation of information flow from V1 to the secondary areas and then on to decoding areas. First, it is possible but unlikely that suppressing activity in cortical areas, as we did, acts on behavior by changing the activity of another area that is truly performing the neural computations for the task. For that to be true, it would require that the visual responses in each of these cortical visual areas are all epiphenomenal. A more likely alternative hypothesis to the sequential interpretation is that multiple brain areas are read out together in parallel. Deficits in visual detection similar to those we observed are also seen with inhibition of the mouse superior colliculus (SC) (***Wang et al., 2020***). Our observations and the SC results suggest that cortical circuits and the colliculus both carry information about visual tasks, and since optogenetic inhibition in each of these studies does not totally abolish the behavior, the areas may each contribute information in near-threshold tasks such as this. This might imply that when animals are performing near perceptual threshold, they use all available representations (e.g. multiple cortical areas, the SC, and potentially others) together to perform the behavior. If it were true that animals do use, or decode, all available representations in sensory areas, this would have implications not just for the circuits used during perceptual tasks, but also for the total amount of information available to the brain about particular stimuli, as calculations of this sort have previously focused only on V1 (***Kafashan et al., 2020; Rumyantsev et al., 2020; Stringer et al., 2019***).

How patterns of neural activity are processed across multiple brain regions to control behavior is a central question about brain function. The role of secondary visual areas in sensory behaviors shown by these results suggests that the mouse visual system, like the visual systems of other species, contains a set of cortical areas that work in concert during perceptual behavior.

## Methods

### Key Resources table

| Reagent type (species) or resource | Designation | Source or reference | Identifiers | Additional information |
| --- | --- | --- | --- | --- |
| Chemical compound | Tamoxifen | Sigma-Aldrich | T5648-5G |  |
| Chemical compound | PEDOT; poly(3,4-ethylenedioxythiophene) | Sigma-Aldrich | 687553 |  |
| Chemical compound | PSS; poly(styrenesulfonate) | Sigma-Aldrich | 243051 |  |
| Genetic reagent | VGAT-ChR2-EYFP | The Jackson Laboratory | RRID:IMSR_JAX:014548 |  |
| Genetic reagent | PV-IRES-Cre | The Jackson Laboratory | RRID:IMSR_JAX:008069 |  |
| Genetic reagent | ReaChR-mCitrine | The Jackson Laboratory | RRIS:IMSR_JAX:024846 |  |
| Genetic reagent | Ai148 | The Jackson Laboratory | RRID:IMSR_Jax:030328 |  |
| Genetic reagent | Cux2-CreERT2 | MMRRC | RRID:MMRRC_032779-MU |  |
| Recombinant DNA | AAV9-hSyn-jGCaMP7s | Addgene | RRID:Addgene_104487 |  |
| Software | MWorks | The MWorks Project |  | mworks.github.io |
| Software | Offline Sorter | Plexon | RRID:SCR_000012 |  |
| Other | C&B Metabond | Parkell | S380 |  |
| Other | Kwik-sil | World Precision Instruments | KWIK-SIL |  |

### Animals

All experimental procedures were approved by the NIH Institutional Animal Care and Use Committee (IACUC) and complied with Public Health Service policy on the humane care and use of laboratory animals. *VGAT-ChR2-EYFP* mice (ChR2 targeted at the *Slc32a1* locus ***Zhao et al., 2011***, Jax stock no. 014548, N=12) were used for behavioral and electrophysiology experiments. Tamoxifen-treated *Ai148;Cux2-CreERT2* mice (N=3, GCaMP6f targeted at the *Igs7* locus and CreERT2 targeted at the *Cux2* locus, respectively***Daigle et al., 2018***; Jax stock No. 030328, MRRC stock no. 032779-MU) were used for imaging experiments in ***Figure 1***–***Figure Supplement 1***. A PV-Cre;ReaChR-mCitrine mouse (Cre targeted at the *Pvalb* locus and ReaChR targeted at the *Gt(ROSA)26Sor* locus ***Lin et al., 2013***; Jax stock nos. 008069 and 024846, respectively) was used for LM inhibition during widefield GCaMP imaging in ***Figure 5***. Eight females and eight males were used in total, singly housed on a reverse light / dark cycle.

### Cranial window implantation and viral injection

Mice were given intraperitoneal dexamethasone (3.2 mg/kg) and anesthetized with isoflurane (1.0-3.0% in 100% O_2_ at 1 L/min). Using aseptic technique, a titanium headpost was affixed using C&B Metabond (Parkell) and a 5 mm diameter craniotomy was made, centered over V1 (−3.1 mm ML, +1.5 mm AP from lambda). A 5 mm cranial window was cemented into the craniotomy, providing chronic access to visual cortex and surrounding higher visual areas.Post-surgery mice were given sub-cutaneous 72-hour slow-release buprenorphine (0.50 mg/kg) and recovered on a heating pad. For experiments involving widefield calcium imaging during LM inhibition, AAV9-hSyn-jGCaMP7s was diluted 1:9 in sterile PBS, and injected 300 um below the surface of the brain. Multiple 800 nL injections were done at 200 nL / min to achieve widespread coverage across the 5 mm window.

### Hemodynamic imaging & visual area map fitting

To determine the location of V1 and surrounding visual areas, we delivered small visual stimuli to head-fixed animals at different retinotopic positions and measured hemodynamic-related changes in absorption by measuring reflected 530 nm light (***Schuett et al., 2002***). Light was delivered with a 530 nm fiber coupled LED (M350F2, Thorlabs) passed through a 530 nm-center bandpass filter (FB530-10, Thorlabs). Images were collected on a Zeiss Discovery stereo microscope with a 1X wide-field objective through a green long-pass emission filter onto a Retiga R3 CCD camera (QImaging, Inc., captured at 2 Hz with 4×4 binning). We presented upward-drifting square wave gratings (2 Hz, 0.1 cycles/degree) masked with a raised cosine window (10° diameter) at different retinotopic locations for 5 seconds with 10 seconds of mean luminance between each trial. Stimuli were presented in random order at six positions in the right monocular field of view. The hemodynamic response to each stimulus was calculated as the change in reflectance of the cortical surface between the baseline period and a response window starting 2-3 s after stimulus onset. This window corresponds to the previously reported timecourse of visually evoked hemodynamic responses in mouse V1 (***Heimel et al., 2007***). Visual area maps were fitted to the cortex based on centroid of each stimulus response.

### Widefield GCaMP Imaging

To validate the placement of the hemodynamic map fits, we applied our hemodynamic imaging of V1 and map fitting procedure in animals in which we also performed GCaMP imaging of V1 and secondary visual areas. Animals (*Ai148;CuX2-CreERT2*; given tamoxifen at P18-22) expressing GCaMP6f in L2/3 cortical excitatory cells were head fixed while a Gabor stimulus (same stimulus used in behavior: +25° azimuth, 0° elevation, 12° in diameter, spatial frequency 0.1 cycle/degree) was shown in their right monocular visual field. We recorded GCaMP fluorescence changes through the 5 mm cranial implant over the contralateral visual cortex. Green fluorescence images were captured using an eGFP filter, at an average frame rate of 10 Hz (1 stimulus frame followed by 32 post-stimulus frames, 4×4 binning) for 150 stimulus presentations. A 10 uL water reward was given randomly (10% probability) to keep animals awake and alert.

For GCaMP imaging with PV-ReaChR inactivation in LM/AL (***Figure 5***; AAV-GCaMP7s), we decomposed the imaging data using principal components analysis (PCA), discarded the first component, which encoded global fluctuation of the entire frame, and reconstructed the image stack. We then took a frame-by-frame difference (*F*_*diff*_) at each pixel to approximate a deconvolution (***Zatka-Haas et al., 2020***). Image response maps show *F*_*diff*_ − *F*_*diff0*_/*F*_0_, where *F*_*diff0*_ is *F*_*diff*_ averaged over 3 frames (300 ms) just before each stimulus onset, and *F*_0_ is average fluorescence over the same baseline frames. Insets in ***Figure 5L,M***, to illustrate decay dynamics, show Δ*F* /*F* ≜ (*F* − *F*_0_)/*F*_0_, where *F* is the pixel value without frame-by-frame differencing and *F*_0_ is as above.

### Behavioral Task

Mice were head-fixed in sound-attenuating custom boxes, and trained to hold a lever and release it when a Gabor stimulus increased its contrast (***Histed et al., 2012***). Mice were required to a lever for a maximum of 4.1 s. At a random time point during this period, a small Gabor patch (+25° azimuth, 0° elevation, 12° in diameter, spatial frequency 0.1 cycle/degree) was presented for 100 ms and animals had 550 ms to report the stimulus by releasing the lever. Possible trial outcomes included a correct release within the reaction time window (hit), a failure to release the lever in response to the stimulus (miss), or a false alarm trial where the animal released the lever before the stimulus had appeared (early). Lever releases occurring in the first 100 ms following the stimulus presentation were counted as earlies or false alarms, as the animal could not have possibly perceived the stimulus and reacted in 100 ms. Trials resulting in hits were rewarded with 1.5-3 uL liquid reward. We began training with a full-screen, 100% contrast stimulus. As animals learned to perform in this task, the size and contrast of the stimulus were decreased and location of the stimulus was slowly moved to the desired azimuth and elevation. As performance stabilized, we also increased the number of contrast levels to cover a range of contrasts or difficulties in order to determine the animals psychometric threshold (the contrast at which the animal was getting 63% of trials correct). Sessions lasted until the animal reached its daily water supplement or until the animal stopped on its own, whichever came first (mean session length = 384 ± 89 trials). Because head-fixed mice make rare eye movements and SC inhibition does not systematically change those movements (***Wang et al., 2020***), mice have no visual streak and small difference in ganglion cell density across the retina, the secondary areas are small enough that our spots covered a large part of their retinotopic maps, and the few eye movements made by head-fixed mice are thought to be compensatory movements that occur with attempted head rotation (***Meyer et al., 2020***), we did not record animals’ eye movements.

### Optogenetic Inhibition

Fiber optic cannulae (400 um diameter 0.39 NA, Thorlabs CFMLC14L02) were dental-cemented above specific areas as determined by hemodynamic intrinsic imaging. Spot size was determined as the full-width half-maximum (FWHM) of the illuminated area (spot area mean ± SEM, 0.42 ± 0.13 mm^2^, N = 30). We used a fiber-coupled LED light source (M470F3, ThorLabs) to deliver illumination with a peak wavelength of 470 nm to the brain. Once animals were performing well on the visual task and we had a full psychometric curve, we introduced a pulsed LED (on for 200 ms, off for 1000 ms) to each trial. The LED pulse train began with the start of the trial and pulsed 0-5 times before the visual stimulus was presented. On 50% of trials, the visual stimulus was presented 55 ms before the last LED pulse in the train (ON trials). On the remaining 50% of trials, the visual stimulus was presented 180 ms before the last LED pulse (OFF trials). Visual onset times were distributed according to a geometric distribution, whose mean was adjusted to balance the desire for a flat hazard function (to discourage waiting for later stimulus times; often reflected in false alarm rates < 5% of trials) and to keep false alarms less than approximately 50% of trials. Once a visual onset time was randomly chosen, it was adjusted by lowering it to either 55 ms after the previous light pulse onset (ON trials) or 820 ms after the previous light pulse offset (OFF trials).

### Behavior Analysis

All analyses were performed with Python. Threshold increases were determined for each behavioral session by fitting a Weibull cumulative density function (hit rates) or Naka-Rushton function (*d*′; ***Herrmann et al., 2012***) across contrast levels. The upper asymptote (lapse rate) and threshold (63% point between the upper and lower asymptotes, Weibull; 50% point, Naka-Rushton) were fit for each trial type with a single slope for LED ON and OFF trials. Only sessions with low lapse rates (<20%) and sufficient trials at each contrast (average 10 trials per contrast level, per trial type) were included for further analysis (539 sessions total; 166 excluded, 373 sessions included). To determine the 100% crossing point for each spot, or the intensity required to double the psychometric threshold, a piecewise-linear function was fit to all percent increases obtained from various LED intensities. Fits were initially done on V1 data with variable slopes. The slopes of spots with low slope CI ratios (< 9 upper:lower CI, ***Figure 1–Figure Supplement 2B***) were averaged and the regressions were re-fit using this determined slope (268.7). Piecewise-linear functions were fit in the same manner described above (***Figure 4–Figure Supplement 2***), first with variable slopes to V1 data and then again with a fixed average slope (261.2), in order to determine a *d*′100% crossing point for each spot. Heatmaps were generated by averaging the 100% crossing point at each pixel. Pixels with no data are black.

### Electrophysiological recordings

For recording experiments, we first affixed a 3D printed ring to the cranial window to retain fluid. With the ring in place, we removed the cranial window and flushed the craniotomy site with sterile normal saline to remove debris. Kwik-Sil silicone adhesive (World Precision Instruments) was used to seal the craniotomy between recording days. Optogenetic light was delivered using the same optical fiber and cannula used for behavioral experiments. The cannula distance from the dura was adjusted to provide a light spot with a full width at half maximum intensity of 0.39 ± 0.19 mm (mean ± SEM, all spots N=13, measured with a USB digital macro documentation microscope, Opti-TekScope). We targeted a multisite silicon probe electrode (NeuroNexus; 32-site model 4×8-100-200-177; 4 shank, 8 sites/shank; sites electrochemically coated with PEDOT:PSS [poly(3,4-ethylenedioxythiophene): poly(styrenesulfonate)]) to the center of the LED spot using a micromanipulator (MPC-200, Sutter Instruments). With the fiber optic cannula and electrode in place, we removed the saline buffer using a sterile absorbent triangle (Electron Microscopy Sciences, Inc.) and allowed the dura to dry for 5 minutes. After insertion, we waited 30-60 minutes without moving the probes to reduce slow drift and provide more stable recordings. We isolated single and multiunit threshold crossings (3 times RMS noise) by amplifying the site signals filtered between 750 Hz and 7.5 Khz (Cerebus, Blackrock microsystems). The visual stimulus (90% contrast Gabor, 100 ms duration, same as during behavioral experiments) was presented on every trial, paired with a 200 ms LED pulse at varying intensity (180 reps each intensity). To keep animals awake and alert, animals were water-scheduled and a 1 uL water reward was randomly provided on 5% of the stimulus trials; we verified animals were licking in response to rewards during the experiments.

### Electrophysiology Analysis

Spike waveforms were extracted and sorted using Offline Sorter (Plexon, Inc.). Single units were identified using waveform clusters that showed separation from noise and unimodal width distributions. Single units had SNR (***Kelly et al., 2007; Histed, 2018; Sanzeni et al., 2020***) of 3.5 ± 0.36 (mean ± std, N = 171 units) and multiunit SNR was 3.1 ± 0.28 (mean ± std, N = 54 units). Multi units were excluded from further analysis. In addition to examining the overall average spike rates of the responses, we calculated mean activity in three response windows: a baseline (r_*base*_) window 40 ms in duration ending 10 ms prior to the delivery of the optogenetic stimulus, an inhibitory minimum (r_*min*_) window of 25 ms starting 25 ms after optogenetic stimulation that ends just before spikes arrive to V1 (∼40 ms, ***Sanzeni et al., 2020***), and a maximum visual response (r_*vis*_) window of 50 ms starting 65 ms post visual stimulus for V1 and lateral areas. The max window was shifted 25 ms later for PM due to delay in the midpoint of the visual response (***Figure 1–Figure Supplement 1, Figure 5–Figure Supplement 1***). Visually responsive units were identified as having an r_*vis*_ 10% or greater over r_*base*_ with no inhibition (0 mW/mm^2^ optogenetic stim). Response traces were baseline corrected to r_*min*_, and the normalized change in spikes rates calculated as r_*vis*_ - r_*min*_ / r_*vis*_ at 0 mW/mm^2^.

## Acknowledgments

We are grateful to Victoria Scott for assistance with animal husbandry and to Patrick Wright for technical advice on hemodynamic imaging. We thank Richard Krauzlis, Bruno Averbeck, and members of the Histed lab for their comments on the manuscript.

---

**Figure 1–Figure supplement 1.**
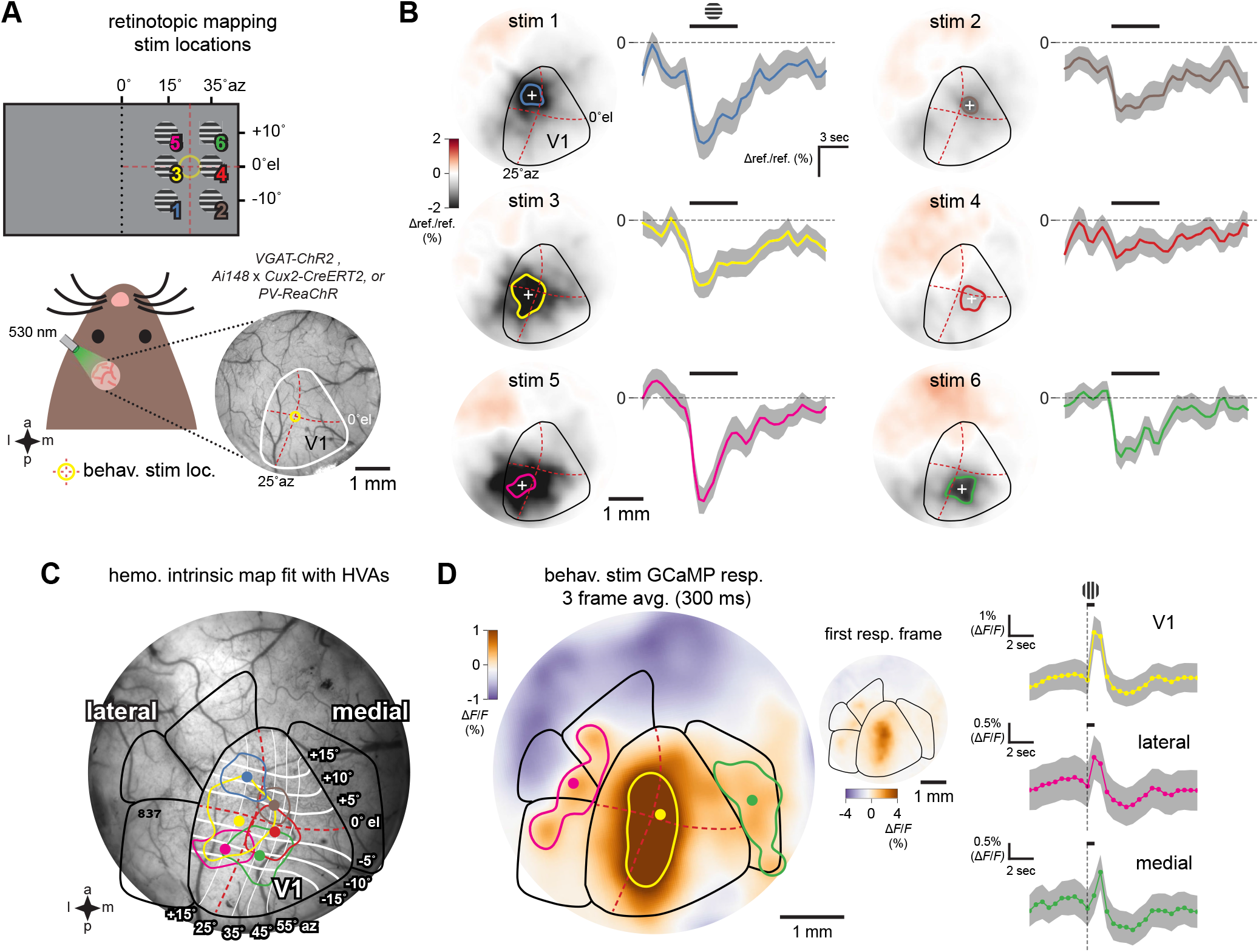
Intrinsic hemodynamic and GCaMP imaging to fit and test visual area maps **(A)** Schematic of hemodynamic intrinsic imaging. Upward drifting Gabor stimuli were presented to head fixed animals randomly at 6 locations in their right visual field (Methods). Each stimulus was shown to the animal 30 times. The cortical surface was evenly illuminated using a fiber coupled 530 nm LED and imaged at 2 Hz through a bandpass green emissions filter. Example behavior visual stimulus location is shown at the center of the hemodynamic map (yellow circle at 25° azimuth (az), 0° elevation (el), red dotted lines). **(B)** Averaged response map and time course of cortical reflectance change at all 6 stimulus locations. Increased cortical blood flow due to the visual stimulus results in a drop in reflectance (Δref./ref.) The period of maximal change in Δref./ref. (%) occurs 2 s after stimulus presentation (***Heimel et al., 2007***). Colored dot and boundary indicates the centroid and 50% contour. **(C)** Scaled and rotated retinotopic map of mouse V1 and secondary visual areas positioned using the 6 hemodynamic response locations in an *Ai148; Cux2-CreERT2* animal which expresses GCaMP6f in cortical excitatory neurons. Elevation and azimuth contours from ***Zhuang et al***. (***2017***). **(D)** GCaMP6f response, averaged over 3 frames, to the visual stimulus used in behavior (a Gabor) presented for 100 ms with 5 s of 50% grey screen between presentations. GCaMP6f fluorescence increases (Δ*F* /*F*) are prominent and well delineated by the area map fit to the hemodynamic responses in V1, and the lateral and medial areas. Colored dot and boundary indicates the centroid and 50% contour. Inset shows single frame GCaMP6f response. Right, time courses of GCaMP6f responses. PM response latency is noticeably longer compared to either V1 or the lateral areas, consistent with electrophysiology results (***Figure 5–Figure Supplement 1***.

**Figure 1–Figure supplement 2.**
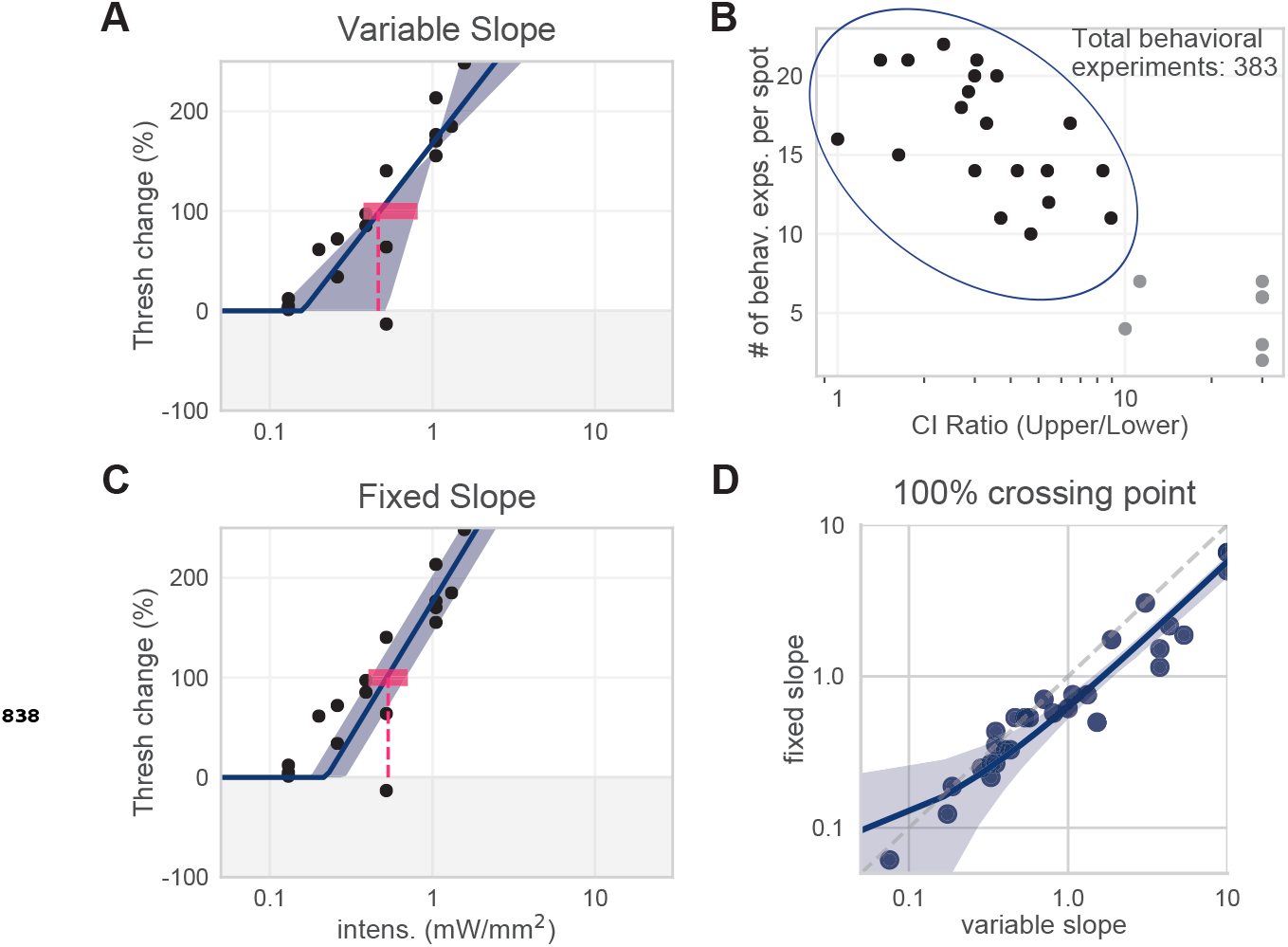
Slope determination for piecewise-linear functions were fit via least-squares regression. All spots from all areas were first fit with a variable-slope piecewise-linear function. We averaged the variable slopes of spots with low slope CI ratios to determine the fixed slope used across all spots from all areas to determine 100% crossing points (266.7 % threshold change / mW/mm^2^). **(A)** Example spot with variable slope (mean 100% crossing point 0.46, bootstrap 95% CI 0.38-0.81 mW/mm^2^). **(B)** Slopes of all spots with low slope CIs (< 9, within blue circle) were averaged to produce a single fixed slope. **(C)** Same example spot as (A), with fixed slope (mean 100% crossing point 0.53, bootstrap 95% CI 0.40-0.66 mW/mm^2^). **(D)** Fixing the slope did not dramatically affect the 100% crossing points. Spots included here are from all visual areas.

**Figure 3–Figure supplement 1.**
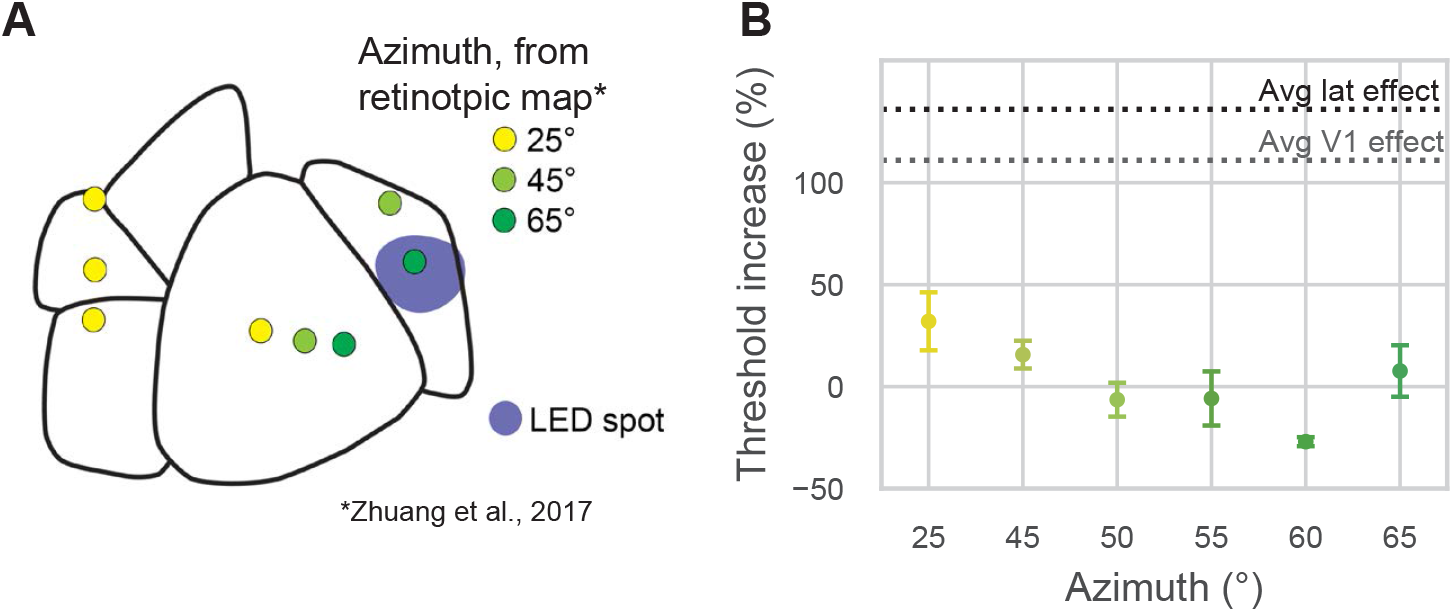
Moving the visual stimulus does not affect PM results. In the retinotopic map of area PM, stimuli at 0° elevation and 25° azimuth are at the very anterior edge of PM. A stimulus at the same elevation but 45° azimuth is in the anterior part of PM but within the PM boundary, and a stimulus at 65° azimuth is closer to the center of PM (***Zhuang et al., 2017***; ***Garrett et al., 2014***). We trained an animal to perform the task with the stimulus at different positions, ranging from 45° to 65° azimuth, in 5° increments. The results were similar to those seen with a 25° azimuth stimulus: little change in threshold when PM was inhibited, confirming that the small PM effects we found were not due to the retinotopic position of the stimulus. **(A)** Map of visual areas, LED spot (blue), and predicted retinotopic locations based on (***Zhuang et al., 2017***; ***Garrett et al., 2014***; yellow = +25° Az, 0° El; light green = +45° az, 0° el; dark green = +65° az, 0° el). **(B)** Average threshold increases (± SEM) at each stimulus location (at 25°, N = 18; all other locations, N = 2 experimental days). Light intensity was held constant at 0.5mW/mm^2^. Average V1 threshold increase at 0.5mW/mm^2^ is shown in light grey (111%). Average LM/AL & RL threshold increase at 0.5mW/mm^2^ is shown in black (136%). Threshold increase at 25° azimuth was 32.0 ± 58.6 (mean ± std. dev). Threshold increase at more lateral azimuths was -3.18 ± 17.4 (mean ± std. dev). At all azimuths tested in PM, the behavioral effects were smaller than those observed when inhibiting V1 or lateral areas LM/AL or RL.

**Figure 4–Figure supplement 1.**
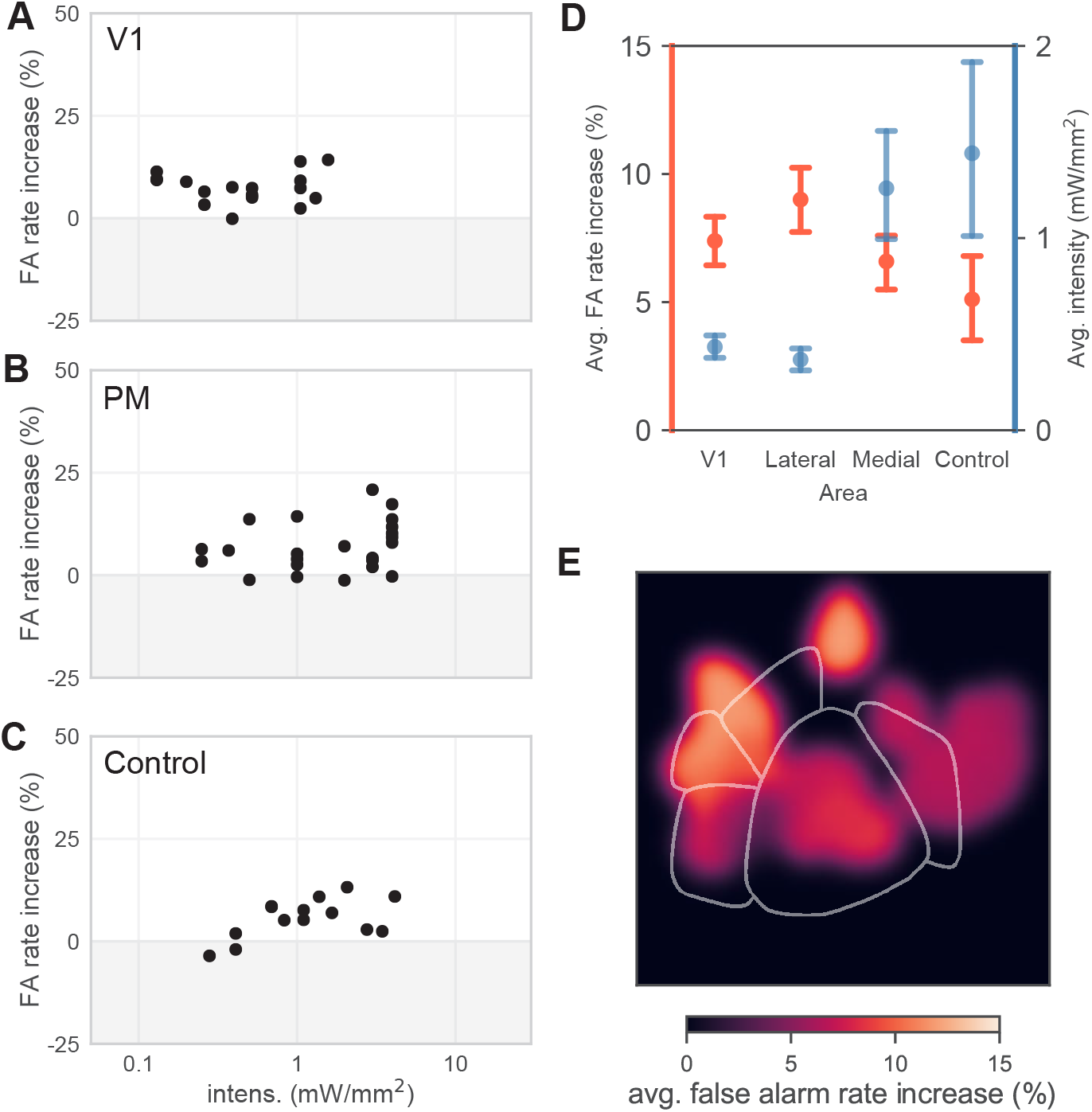
False-alarm rates do not show systematic variation across areas. **(A-C)** Changes in false-alarm rates between ON and OFF trial types show little correlation with light intensity. Therefore, it would be inappropriate to predict a crossing point in the same manner as we did for hit-rate and *d*′. False-alarm rate analysis below is done using average false-alarm rates across all intensities. **(D)** We saw little significant difference across areas in average false-alarm rate changes (orange). To rule out that different light intensities used in different areas could contribute to changes, we computed the average intensity used in each area (blue). False alarm rate changes range between 5-10% in all four areas, with a small trend for smaller FA changes in medial and control areas. Higher average intensities were used in medial areas and control areas, as sensitivity changes were not seen at low intensities. **(E)** Average false-alarm rate map generated by averaging the false-alarm rate increase at each pixel. False-alarm rates were generally small (0-10%). Pixels with no data are black. For consequences of false alarm rate (and hit rate) variation on perceptual performance, see comparison of hit-rate and *d*′thresholds (Results, **Figure 1, Figure 4** and ***Figure 4–Figure Supplement 2***).

**Figure 4–Figure supplement 2.**
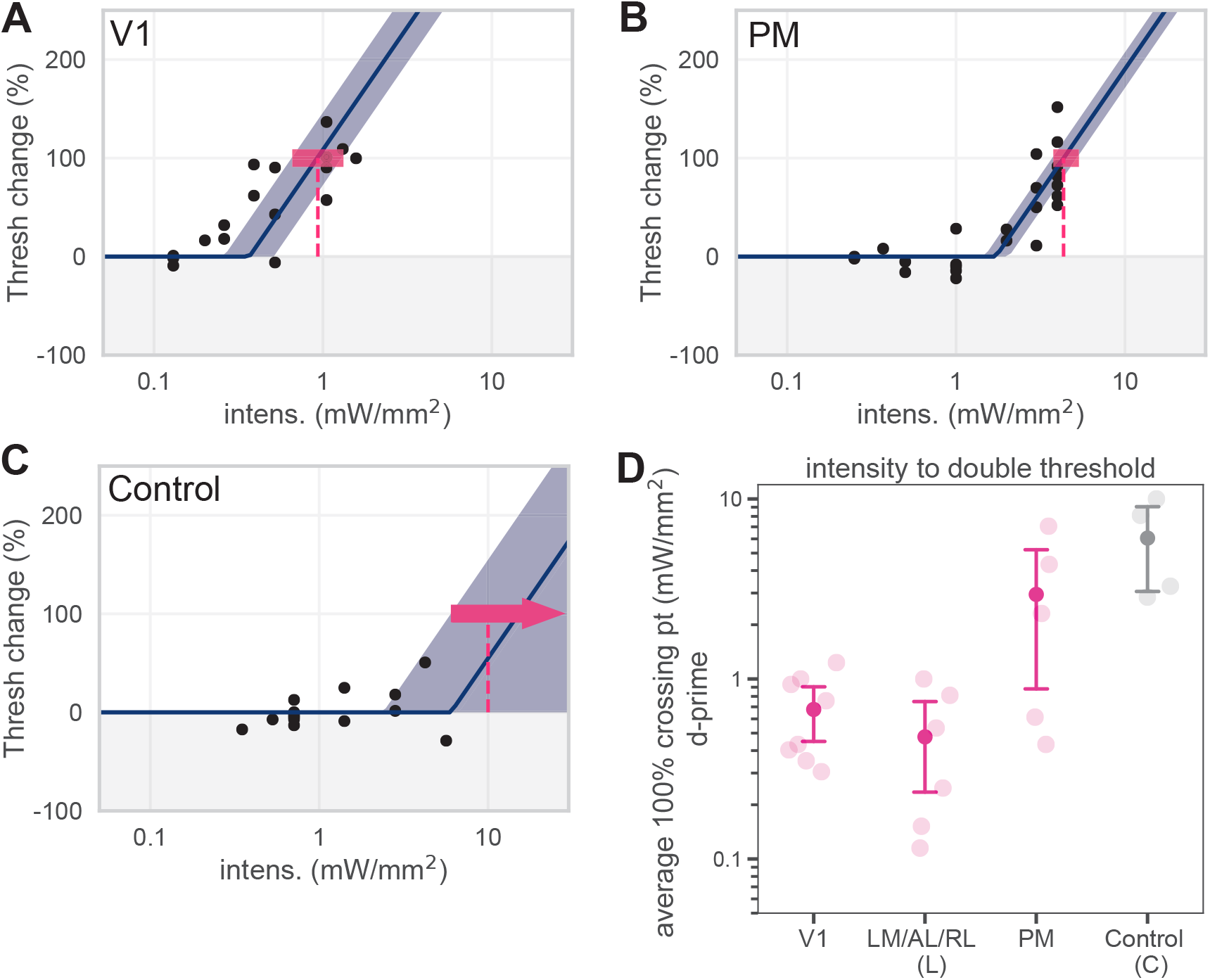
Predicting 100% crossing point using *d*′values does not alter behavioral effects. **(A-C)** Piecewise-linear functions were fit to *d*′data in the same manner described in ***Figure 1–Figure Supplement 2*** to predict the intensity necessary to double sensitivity (the *d*′100% crossing point). **(D)** Plotting behavioral effects in terms of sensitivity (*d*′) does not affect our main conclusion that inhibiting lateral areas (LM/AL/RL) produce effects similar to those seen in V1, while inhibiting PM produces effects similar to those seen in control areas.

**Figure 4–Figure supplement 3.**
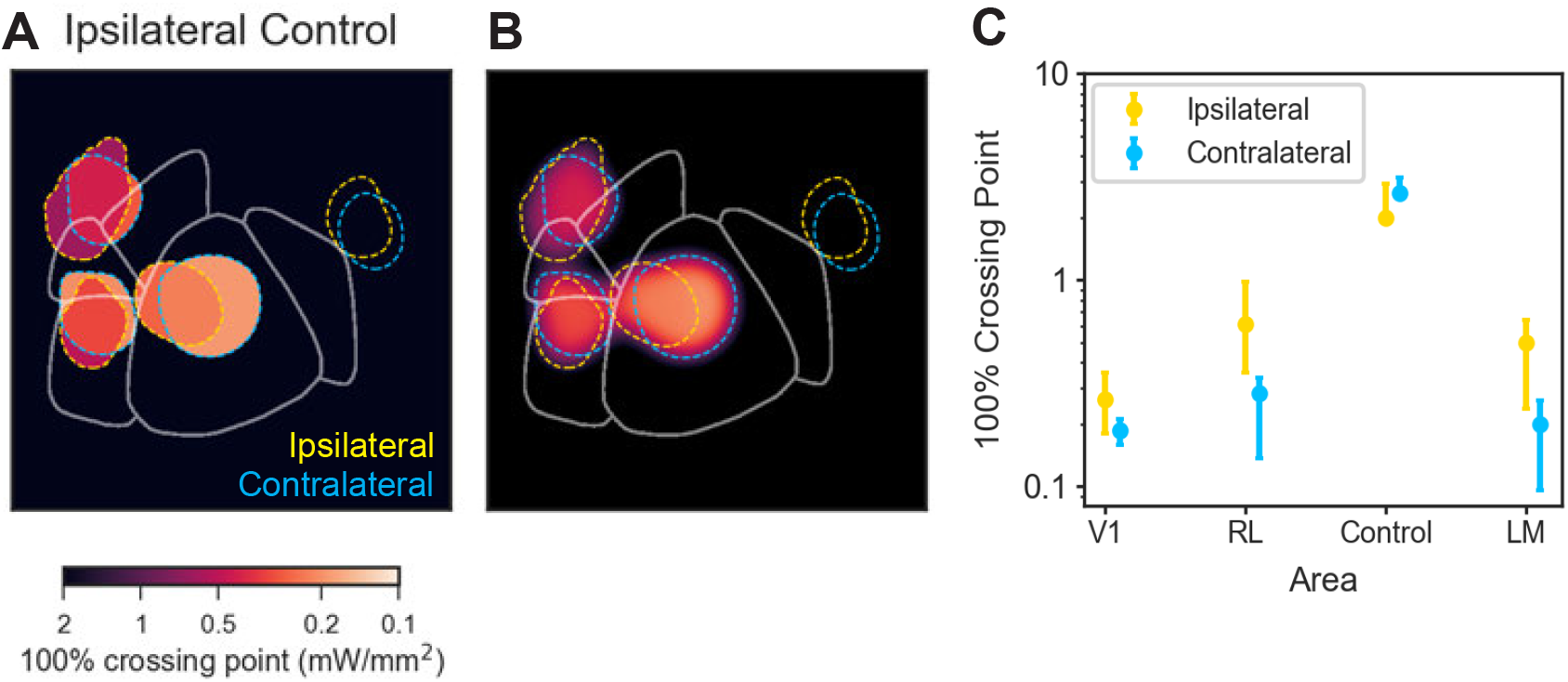
Changing the motor response from contralateral to ipsilateral paw produced a similar pattern of behavioral changes. Thus, observed behavioral deficits are primarily sensory, not motor, effects. **(A-B)** In all other data reported in this work, animals were trained to use the paw contralateral to the inhibited hemisphere to press and release the lever. To determine how effects would change when the motor response modality was changed, we trained one animal to use its ipsilateral paw and inhibited areas matched to those inhibited in an second animal trained to use its contralateral paw. Inhibiting similar areas in the two animals (yellow dashed lines surround spots from animal using its ipsilateral paw, blue dashed lines: contralateral paw) produced similar results. **(C)** Crossing points (with 95% CIs for the piecewise-linear functions) for the data shown in A-B. There is a trend for the ipsilateral-paw animal to have a mean crossing point larger (*i.e*. require more power to achieve a similar change in threshold) than the contralateral-paw animal for areas V1, RL, LM/AL. However, in none of these cases do the thresholds do not rise to the level of the control data. Therefore, while we cannot rule out that there may be some motor contribution to the inhibition-induced behavioral changes, changing the animals’ motor response shows that the effects on performance are primarily non-motor (sensory) effects.

**Figure 5–Figure supplement 1.**
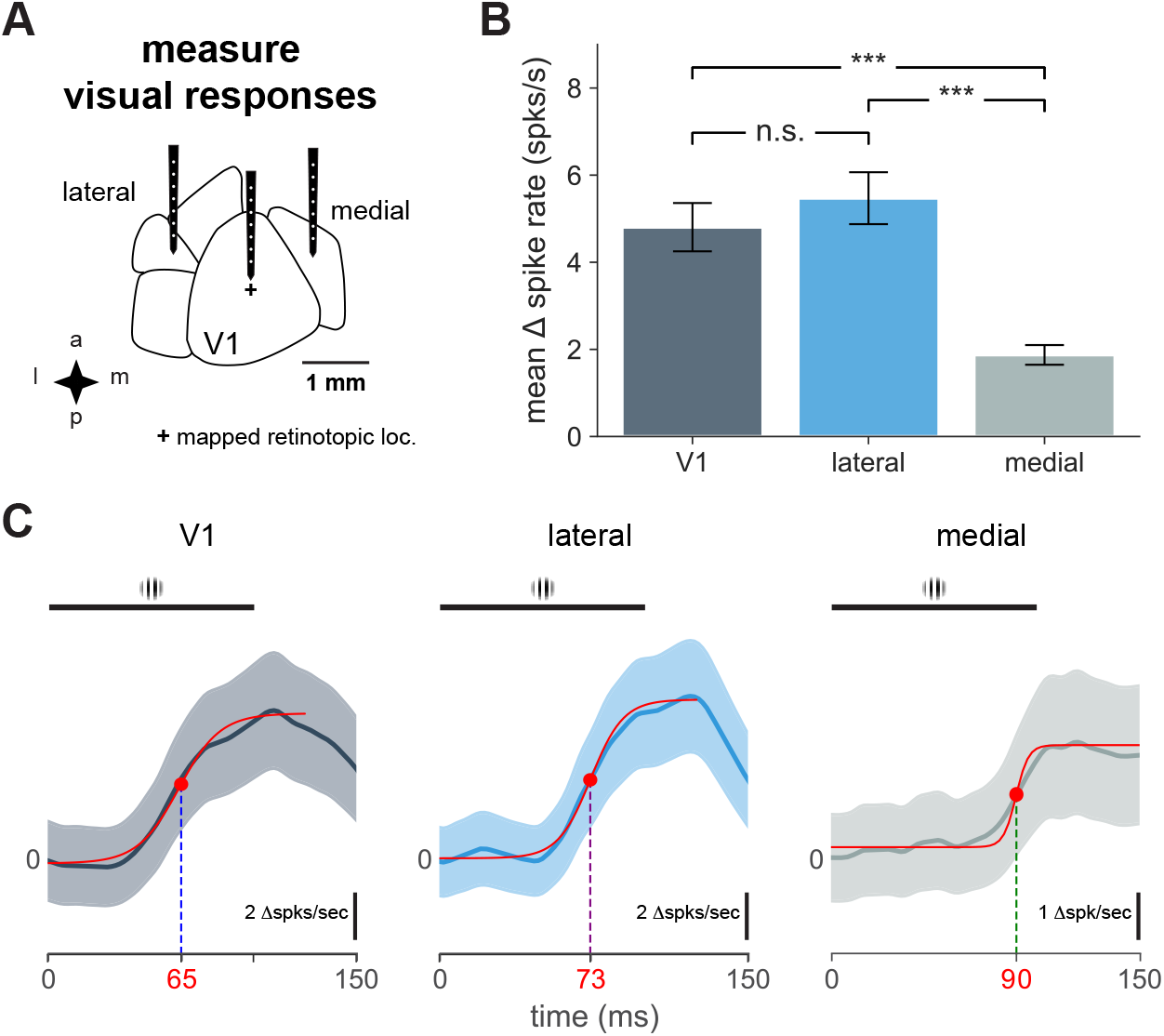
Characteristics of V1 and secondary visual area cortical responses to visual stimulus and response timing. **(A)** Schematic of electrophysiological recordings in V1, lateral, and medial areas. Plus sign indicates visual stimulus at the retinotopic location of the stimulus (identified with hemodynamic imaging) in V1. **(B)** Mean change in spike rate recorded in indicated cortical area in response to 100 ms flashed Gabor. (mean ± SEM: V1, N = 14 units; lateral, N = 10 units; medial, N = 16 units; t-tests: V1-lateral, n.s, *t* = 0.80, *p* = 0.49; V1-medial, ***, *t* = 5.13, *p <* 0.0001; lateral-medial, ***, *t* = 6.57, *p <* 0.0001). **(C)** Plot of average unit responses (visual stimulus presentation at time = 0) for indicated cortical area with sigmodial fit (solid red lines). Black bar indicates visual stimulus. Time point for half-maximal response shown for each fit with red dot.

**Figure 5–Figure supplement 2.**
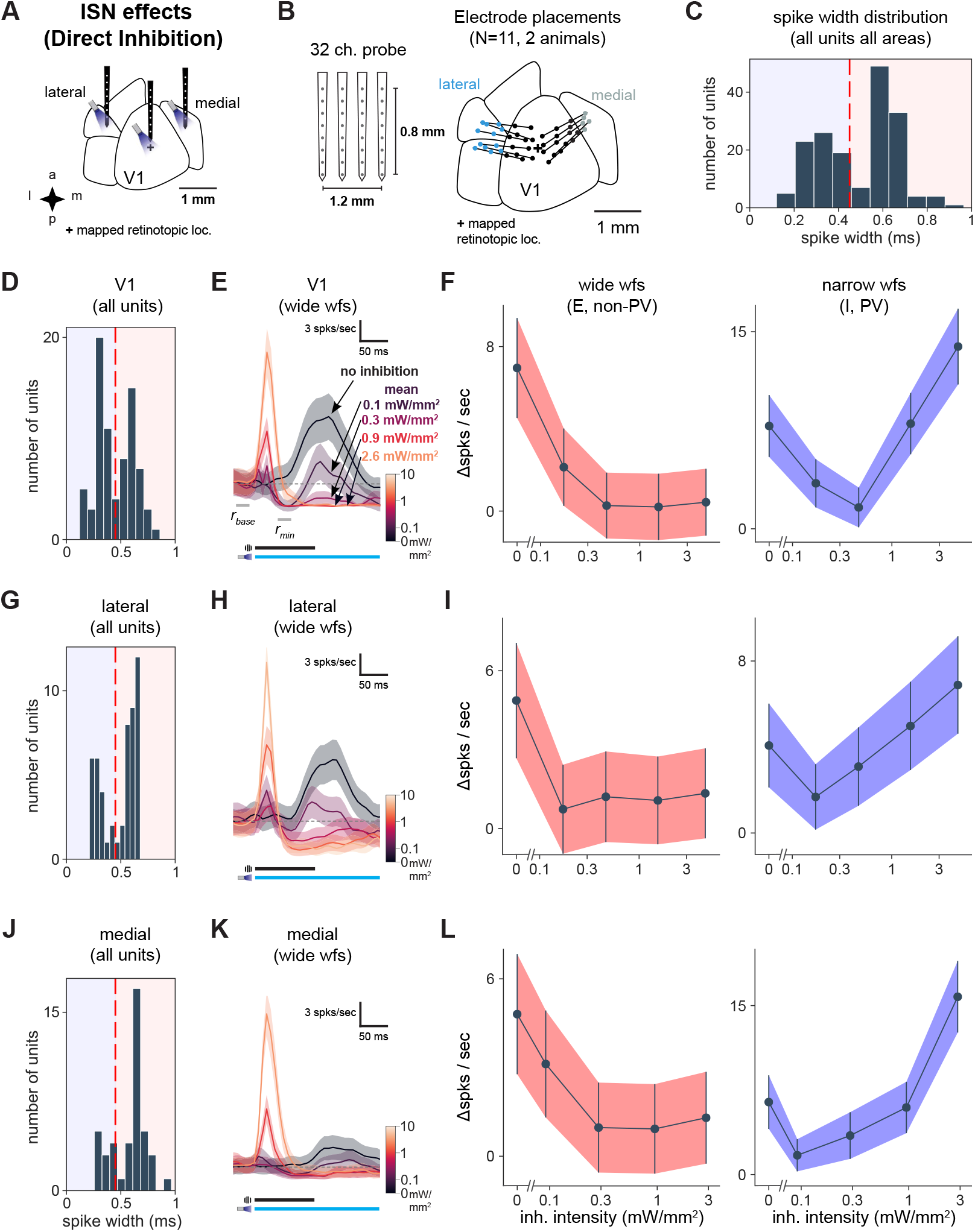
V1, lateral areas, and PM display characteristics of inhibition stabilization. **(A)** Schematic of direct optogenetic inhibition of V1, lateral areas, and PM to examine ISN-related effects. Plus sign indicates V1 retinotopic location of the visual stimulus, identified with hemodynamic intrinsic imaging (Methods). **(B)** Schematic of 4 recording shanks and electrode placements superimposed on cortical area map. Dots: individual shanks. Lines connect shanks recorded in a single experiment. Color: cortical area. **(C)** Spike width histogram of all units across all areas recorded, showing bi-modal distribution, N = 171 units). We used 450 ms as a cutoff between wide and narrow units (red dotted line). The narrow-waveform units reflect mainly inhibitory units, likely dominated by PV+ fast-spiking cells, while wide-waveform units are majority excitatory cells and also some inhibitory neurons (***Sanzeni et al., 2020***). **(D)** Spike width histogram of all V1 units (N = 77). **(E)** Average response of wide-waveform (E) visually responsive V1 units (N = 10) to optogenetic light pulses combined with a high-contrast visual stimulus. Initial transient is due to inhibitory neurons’ early spikes and rapidly decays. **(F)** Average responses before visual response (*i.e. r*_*min*_ − *r*_*base*_ in panel E) of wide (left, light red) and narrow (right, blue) waveform units (wide, N = 41 units; narrow, N = 36 units). As expected, wide-waveform (E plus some I) units show little increase at the highest optogenetic power, while narrow-waveform (I) units first decrease, and then increase, their firing rates as inhibitory stimulation intensity is increased. **(G)** Spike width histogram of all units from lateral sites (N = 51). **(H)** Average responses of wide-waveform visually responsive lateral units (N = 10 units). Conventions as in E. **(I)** Optogenetic responses of wide- (N = 32) and narrow-waveform (N = 19) units in lateral areas. Conventions as in F. **(J-L)** As for D-F, but for recordings from medial areas. N = 32 wide-waveform, N = 11 narrow-waveform units.

**Figure 5–Figure supplement 3.**
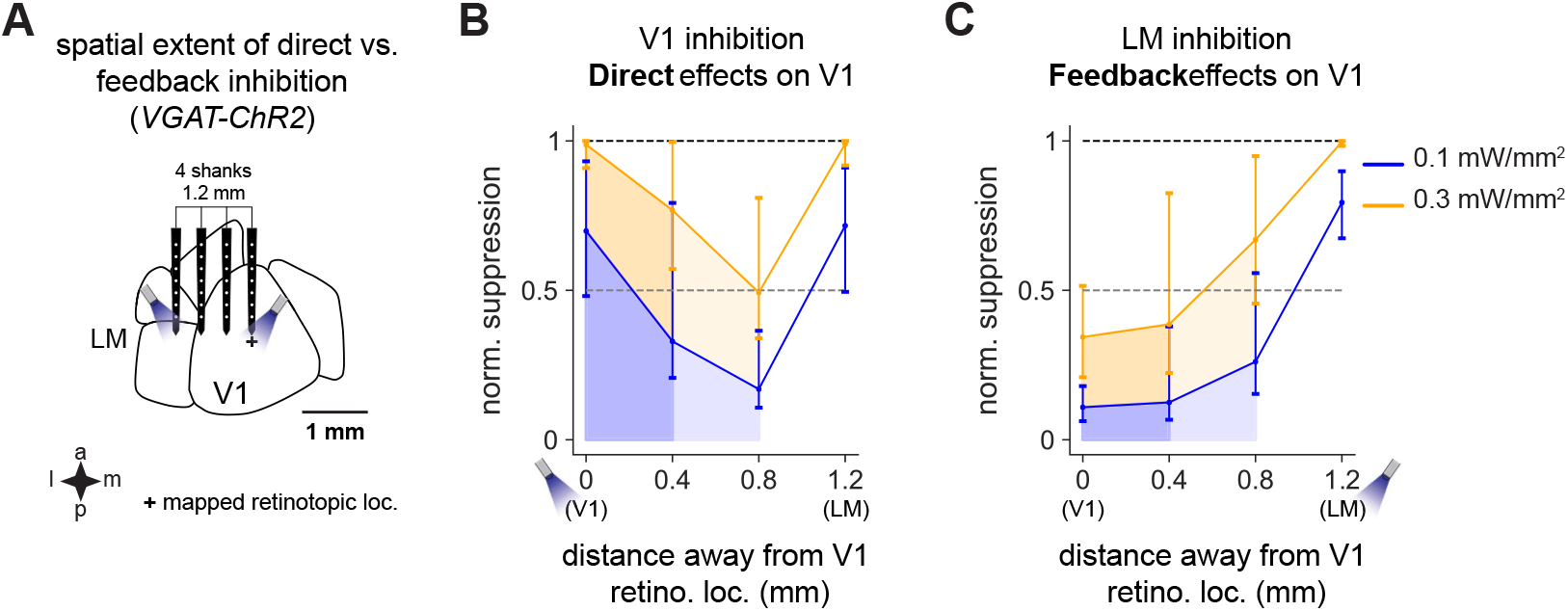
V1 activity is suppressed substantially more when V1 is directly inhibited than via feedback when LM is inhibited. **(A)** Schematic of recording setup. A probe with four shanks was placed across V1 and LM. Light was delivered to V1 or LM to cause inhibition, and responses were recorded across all four shanks. **(B)** Responses in V1 to direct inhibition of V1, showing total normalized suppression (i.e. fraction of the visual response without light that is removed by inhibition; Methods), at distances of 0.4 mm (dark shades) and 0.8 mm (lighter shades) from the recording shank placed at the center of the V1 retinotopic location of the stimulus. Responses are shown at two powers, 0.1 mW/mm^2^ (blue) and 0.3 mW/mm^2^ (orange). Responses are normalized to the suppression at the light location (direct suppression) at 0.3 mW/mm^2^. Normalized suppression increases again at 1.2 mm as the probe crosses into LM and activity is suppressed through inhibition of feedforward connections from V1. **(C)** Feedback effects on V1 caused by LM inhibition. Same conventions as Panel B. At all intensities and at all distances, feedback inhibition causes less V1 suppression overall than direct suppression of V1 (i.e. shaded regions in C versus shaded regions in B).

